# RICTOR Drives ZFX-mediated Ganglioside Biosynthesis to Promote Breast Cancer Progression

**DOI:** 10.1101/2022.01.10.475595

**Authors:** Kajal Rajput, Mohd. Nafees Ansari, Somesh K. Jha, Nihal Medatwal, Pankaj Sharma, Sudeshna Datta, Animesh Kar, Trishna Pani, Kaushavi Cholke, Kajal Rana, Ali Khan, Geetashree Mukherjee, SVS Deo, Jyothi S Prabhu, Arnab Mukhopadhyay, Avinash Bajaj, Ujjaini Dasgupta

**Author notes:** These authors contributed equally to the work (Kajal Rajput, Mohd. Nafees Ansari). Department of Systems Pharmacology and Translational Therapeutics, University of Pennsylvania, Philadelphia, PA, USA. Sudeshna Datta, Data analyst, Institute of Life Sciences, Nalco square, Bhubaneswar-751023, India.

## Abstract

Sphingolipid and ganglioside metabolic pathways are crucial components of cell signalling, having established roles in tumor cell proliferation, invasion, and migration. However, regulatory mechanisms controlling sphingolipid and ganglioside biosynthesis in mammalian cells is less known. Here, we show that RICTOR, the regulatory subunit of mTORC2, regulates the synthesis of sphingolipids and gangliosides in luminal breast cancer-specific MCF-7 and BT-474 cells through transcriptional and epigenetic mechanisms. RICTOR regulates glucosylceramide levels by modulating the expression of UDP-Glucose Ceramide Glucosyl transferase (UGCG). We identify Zinc Finger protein X-linked (ZFX) as a RICTOR-responsive transcription factor whose recruitment to the *UGCG* promoter is regulated by DNA methyltransferases and histone demethylase (KDM5A) that are known AKT substrates. We further demonstrate that RICTOR regulates the synthesis of GD3 gangliosides through ZFX and UGCG, and triggers the activation of EGFR signalling pathway, thereby promoting tumor growth. In line with our findings in cell culture and mice models, we observe an elevated expression of RICTOR, ZFX, and UGCG in Indian luminal breast cancer tissues, and in TCGA and METABRIC datasets. Together, we establish a key regulatory circuit, RICTOR-AKT-ZFX-UGCG-Ganglioside-EGFR-AKT, and elucidate its contribution to breast cancer progression.

## INTRODUCTION

Sphingolipids and gangliosides are metabolically interconnected structural and signalling components of cell membranes, and deregulation in their metabolism is linked to key human diseases including cancer (1, 2). Ceramides are the central hub of sphingolipid pathway, and modifications at their 3’-hydroxyl terminal lead to structurally diverse classes of sphingolipids like glucosylceramides, sphingomyelins, and ceramide-1-phosphates with a distinct role in different facets of tumorigenesis (Figure 1A) (3). Synthesis of glucosylceramides from ceramides is catalysed by UDP-Glucose Ceramide Glucosyl transferase (UGCG), and ceramide-glucosylceramide rheostat connecting sphingolipid and ganglioside metabolism plays a crucial role in tumor progression, drug resistance, and chemotherapeutic response (Figure 1A) (1). Gangliosides are sialic acid-containing glycosphingolipids residing in glycolipid-enriched microdomains (GEMs) of plasma membrane (4). Gangliosides, as residents of GEMs, can induce activation/deactivation of Receptor-Tyrosine Kinases (RTKs), thereby regulating the downstream signalling processes (5). Therefore, understanding the mechanisms that regulate the ceramide-glucosylceramide rheostat connecting sphingolipid and ganglioside pathways, and its impact on RTK signalling may potentially lead to identification of new therapeutic targets.

PI3K/AKT/mTOR is the key downstream pathway of most RTKs (6), and is hyperactivated in 60% of breast cancer patients due to increased expression of growth factors and their receptors or genetic alterations in *PIK3CA, AKT*, and *PTEN* (7). Mammalian-target of Rapamycin (mTOR), an intracellular serine/threonine kinase of PI3K pathway exists in two different complexes, mTORC1 and mTORC2. Rapamycin-Insensitive Companion of mTOR (RICTOR) is the key regulatory subunit of mTORC2 that can directly phosphorylate AKT at Ser^473^ (8). Apart from AKT activation, mTORC2 also regulates tumorigenesis through activation of other substrates like AGC kinases, Serum- and Glucocorticoid-induced protein kinase 1 (SGK), and Protein Kinase C (PKC) (9–12). Amplification of *RICTOR* or elevated RICTOR protein expression is positively correlated with poor overall survival of cancer patients (13–17). Meta-analysis of cancer genomics datasets show co-occurrence of RTK alterations with RICTOR overexpression in different cancer types (18). Therefore, there is a need to unravel the role of RICTOR (mTORC2)-mediated regulatory pathways in tumorigenesis.

Metabolic reprogramming is one of the hallmarks of tumorigenesis, and mTORC2 has emerged as a key link between RTK signalling and cancer metabolic reprogramming (19). mTORC2 can regulate glycolytic metabolism in cancer cells either through AKT phosphorylation, or via regulating c-Myc expression, or through transcription factor, FoxO (19). mTORC2 also regulates lipogenesis, lipid homeostasis, and adipogenesis in non-cancer cells, however, mTORC2-mediated regulation of sphingolipids and gangliosides in cancer cells is not well understood (20). Yeast studies have shown that TORC2-dependent protein kinase, Ypk1, activates phosphorylation of subunits of ceramide synthase complex, leading to an increased synthesis of sphingolipids (21). It is also known that membrane stress in *S. cerevisiae* causes redistribution of Slm proteins in plasma membrane, leading to their association with TORC2 complex, and further activation of sphingolipid synthesis (22). Recent studies have shown that mTORC2 may promote tumorigenesis in hepatocarcinoma by promoting *de novo* fatty acid and lipid synthesis (23). However, mTORC2-mediated regulation of sphingolipid and ganglioside metabolism through transcriptional and epigenetic regulatory mechanisms, and impact of the network surrounding this metabolic hub on tumorigenesis is not known. Herein, we delineate mTORC2-mediated regulation of ceramide-glucosylceramide rheostat, and its role in breast cancer progression.

Luminal (A and B) is most common breast cancer subtype, accounting for ∼75% of all cases (24, 25). Although luminal breast cancer patients, characterized by presence of estrogen and progesterone receptor (ER and PR), have a better prognosis, but recurrence and resistance to endocrine therapy are major challenges (25). Here, we show that RICTOR-mediated epigenomic alterations in DNA and histone methylation lead to rewiring of *UGCG* transcriptional competence in luminal breast cancer (MCF-7 and BT-474) cells. We identify ZFX as the RICTOR-responsive transcription factor that regulates UGCG expression. We further demonstrate that elevated expression of ZFX/UGCG alters ganglioside levels, and activate epidermal growth factor receptor (EGFR)-mediated PI3K/AKT/mTOR/MAPK signalling leading to tumor progression. Finally, Indian patient tumor tissues with high RICTOR expression also manifest high UGCG and ZFX expression, emphasizing the role of mTORC2-regulated ganglioside metabolism in tumor development.

## RESULTS

### Luminal Tumors have High Glucosylceramide Levels and High RICTOR Expression

Ceramide-glucosylceramide rheostat plays a crucial role in tumor progression, drug resistance, and chemotherapeutic response where synthesis of glucosylceramides from ceramides is catalysed by UGCG, and glucosylceramidase (GBA) hydrolyses glucosylceramides to ceramides (Figure 1A, 1B). To detect any imbalance in ceramide-glucosylceramide rheostat in luminal breast cancer patients, we quantified levels of ceramides and glucosylceramides in luminal patient tumor tissues from an Indian cohort (**Dataset 1**), and compared them with adjacent normal tissues using liquid-chromatography mass spectrometry (LC-MS/MS) in multiple reaction monitoring (MRM) mode (26, 27). Tumor tissues showed 1.6-28 fold increase in levels of all ceramide species as compared to their adjacent normal tissues (**Supplementary Figure S1A**). Similarly, we observed a 1.8-100 fold increase in levels of all glucosylceramide species in tumor tissues (**Supplementary Figure S1B**). Glucosylceramide to ceramide ratio for each species is 1.7-42 fold higher in tumor tissues than in normal tissues suggesting that the rheostat is shifted towards glucosylceramides in luminal breast cancer patients (Figure 1C).

To decipher the effect of increased glucosylceramides on tumor progression, we overexpressed UGCG transcript-encoding cDNA in luminal breast cancer representing MCF-7 cells (MCF-7_UGCG^OE^), and used only vector-transfected cells (MCF-7_VECT^OE^) as control. We also silenced UGCG expression by siRNA (MCF-7_UGCG^SL^), and used scrambled siRNA-transfected cells as control (MCF-7_SCRAM^SL^). Overexpression and downregulation of UGCG were confirmed by qRT-PCR (**Supplementary Figure S1C**), immunoblot (Figure 1D, 1E), and immunofluorescence staining (**Supplementary Figure S1D**). As expected, overexpression of UGCG attenuated the levels of ceramides (Figure 1F), and MCF-7_UGCG^OE^ cells showed a 4-12 fold increase in levels of all glucosylceramide species (Figure 1F) that was also validated by immunostaining of ceramides (Figure 1G). MCF-7_UGCG^OE^ cells exhibited elevated cell proliferation as compared to MCF-7_VECT^OE^ cells (Figure 1H). In contrast, UGCG silencing showed a >1.7-fold (*p* < 0.001) decrease in cell proliferation as compared to MCF-7_SCRAM^SL^ cells (Figure 1I). MCF-7_UGCG^OE^ cells also possess enhanced cell migration (**Supplementary Figure S1E**) whereas MCF-7_UGCG^SL^ possess reduced cell migration (**Supplementary Figure S1F**). Similarly, overexpression of UGCG enhanced the cell proliferation of BT-474 cells (**Supplementary Figure S1G**) and UGCG silencing inhibited cell proliferation (**Supplementary Figure S1H**). Mice xenograft studies recorded a significant increase in tumor growth kinetics of MCF-7_UGCG^OE^ tumors (Figure 1J, 1K). We then investigated the effect of UGCG overexpression on RTK signaling pathway to decipher the mechanism of UGCG-mediated enhanced cell proliferation. Immunoblot studies revealed enhanced expression of RICTOR, pAKT^S473^, and pSGK^S78^ in MCF-7_UGCG^OE^ cells (Figure 1L), whereas there was no significant change in RAPTOR and mTORC1 targets p4EBP1 and p70S6K (Figure 1L). We also generated *RICTOR* silenced MCF-7_RICTOR^SH^ cells using lentiviral-mediated shRNA transfection that showed reduced expression of pAKT^S473^ and pSGK^S78^ (Figure 1L). The activation of PI3K/AKT/mTOR pathway on UGCG expression hinted at a complex feed forward loop involving sphingolipid metabolism, RTKs, and PI3K/AKT/mTOR signaling.

RICTOR being a key component of mTORC2, earlier studies on analysis of The Cancer Genome Atlas (TCGA) curated invasive breast carcinoma patient datasets have shown significant correlation of *RICTOR* upregulation, mutation, or amplification with low overall survival (28–30). To validate any deregulation of RICTOR expression in luminal breast tumors, we quantitated the RICTOR expression by immunofluorescence in tumor and adjoining matched normal breast tissues from luminal patients from the same Indian cohort (Figure 1M**, Supplementary Figure S1I**, **Dataset 1**). We observed an abundant (n = 7, *p* = 0.053) increase in RICTOR expression in tumor tissues as compared to adjoining normal tissues (Figure 1N). Above results show that luminal breast cancer tissues have high levels of glucosylceramides and high RICTOR expression. Therefore, we hypothesized that RICTOR (mTORC2) might regulate the sphingolipid metabolism, and increased levels of glucosylceramides may be instrumental for enhanced cancer cell proliferation and tumor progression in luminal breast cancer patients (Figure 1O).

**Figure 1.**
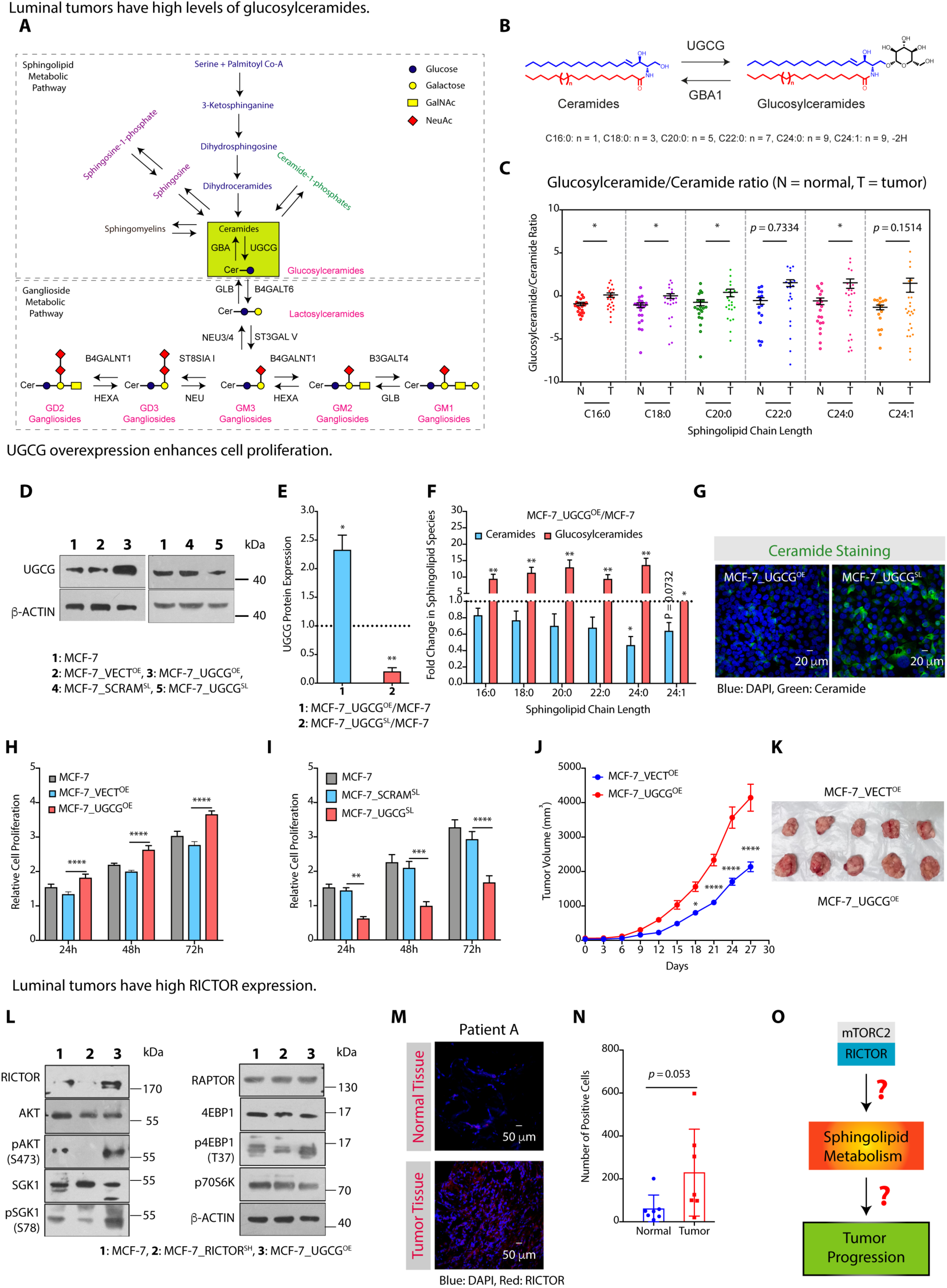
Luminal Tumors have Elevated Glucosylceramide Levels and High RICTOR Expression. **(A)** A schematic presentation showing sphingolipid and ganglioside metabolic pathways along with ceramide-glucosylceramide rheostat connecting these pathways. **(B)** A schematic showing the synthesis of glucosylceramides, with different fatty acyl chain lengths from ceramides catalysed by UGCG, and breakdown of glucosylceramides to ceramides by GBA1. **(C)** Glucosylceramide to ceramide ratios for luminal tumor tissues (labelled as T) and adjacent normal tissues (labelled as N) indicate that the balance is shifted towards the synthesis of glucosylceramides. **(D, E)** Immunoblot with representative β-actin as control **(D)** and quantification (mean ± SD, n = 3) **(E)** validations showing increase in UGCG expression in MCF-7_UGCG^OE^ cells and effective silencing of UGCG in MCF-7_UGCG^SL^ cells. **(F)** Fold change in levels of ceramides (mean ± SD, n = 5) and glucosylceramides (mean ± SD, n = 5) confirm decrease in ceramide levels and increase in glucosylceramide levels in MCF-7_UGCG^OE^ cells over MCF-7 cells. **(G)** Immunofluorescence images using anti-ceramide antibody show low ceramide levels in MCF-7_UGCG^OE^ cells and high ceramide levels in MCF-7_UGCG^SL^ cells. **(H, I)** Cell proliferation (mean ± SD, n = 5) assay demonstrate an increase in proliferation of MCF-7_UGCG^OE^ cells **(H)** and decrease in MCF-7_UGCG^SL^ cells over MCF-7_SCRAM^SL^ cells **(I)**. **(J, K)** Tumor growth kinetics **(J)** and images of the excised tumors **(K)** reveal enhanced growth of MCF-7_UGCG^OE^ tumors as compared to MCF-7_VECT^OE^ tumors (mean ± SEM, n = 5). **(L)** Immunoblots with representative β-actin as control showing expression of RICTOR, RAPTOR and their downstream targets pAKT^Ser473^, SGK1, pSGK1^Ser78^, 4EBP1, p4EBP1^T37^, p70S6K in MCF-7_UGCG^OE^ cells along with representative β-actin as control. MCF-7_RICTOR^SH^ cells are used as a control. **(M, N)** Immunofluorescence images **(M)** and quantification **(N)** (mean ± SD, n = 7) show elevated RICTOR expression in tumor tissues as compared to adjacent normal tissue sections. **(O)** A schematic diagram showing questions that needed to be unravelled in order to understand the mTORC2-mediated regulation of sphingolipid-metabolic pathway and their role in tumor progression. Details of all Immunoblots with originals are provided in supplementary information. Data in Figure 1C, 1E, and 1F were analysed using paired student’s *t* -test. Figure 1H-J was analysed using Two-way ANOVA *p*-value: \**p* < 0.05, \*\**p* < 0.01, \*\*\**p* < 0.001, \*\*\**p* < 0.0005.

### RICTOR Silencing Inhibits Cell Proliferation and Migration

As luminal tumor tissues show high RICTOR expression, we first determined the effect of RICTOR silencing on cancer cell proliferation, migration, and invasion in breast cancer luminal subtype representative MCF-7 and BT-474 cells. We used engineered *RICTOR* silenced MCF-7_RICTOR^SH^ cells and MCF-7_SCRAM^SH^ cells with scrambled shRNA as control (Figure 2A**, Supplementary Figure S2A**). Cellular assay showed ∼1.25-fold (*p* < 0.0001) decrease in proliferation of MCF-7_RICTOR^SH^ cells after 72h (Figure 2B), and >2.0-fold (*p* < 0.0001) decrease in number of colonies during anchorage-dependent assay as compared to MCF-7_SCRAM^SH^ cells (**Supplementary Figure S2B**). Similarly, scratch wound and transwell migration assays showed >1.3-fold (*p <* 0.0001) (**Supplementary Figure S2C**), and >1.5-fold (*p <* 0.001) (**Supplementary Figure S2D**) decrease in number of migrating cells on RICTOR silencing. Similarly, RICTOR silencing inhibited cell proliferation in BT-474 cells (**Supplementary Figure S2E, S2F**). The effect of RICTOR silencing in MCF-7 cells was further validated by quantifying the expression of downstream effectors of mTORC2 pathway including AGC kinases like AKT and SGK. As expected, there was a ∼2-fold decrease in expression of pAKT^S473^ (*p* = 0.09) and pSGK^S78^ (*p* < 0.05) upon RICTOR silencing, and there was no change in expression of mTORC1-specific RAPTOR, p4EBP1^T37^, and p70S6K (Figure 2C, 2D). These experiments suggest that pathogenic activation of RICTOR seen in luminal cancer tissues may activate proliferation through activation of downstream effectors like AGC kinases. Next, we ventured to understand how RICTOR activation may lead to enhanced proliferation, particularly in the context of sphingolipid metabolism.

### mTORC2 Alters Ceramide-Glucosylceramide Rheostat via Regulating UGCG

To decipher the effect of RICTOR silencing on sphingolipid metabolism, we performed quantitative profiling of sphingolipids in MCF-7 and MCF-7_RICTOR^SH^ cells as well as in BT-474 and BT-474_RICTOR^SH^ cells by LC-MS/MS. All species (C16:0, C18:0, C20:0, C22:0, C24:0, and C24:1) of glucosylceramides showed a significant decrease in MCF-7_RICTOR^SH^ cells with a concurrent increase in levels of ceramides as compared to MCF-7 cells (Figure 2E). There is a 2-9 fold increase in ceramide levels, and 1.3-1.7 fold decrease in glucosylceramide levels in MCF-7_RICTOR^SH^ cells over MCF-7 cells (Figure 2F, G). Immunofluorescent staining with anti-ceramide antibody elicited an increase in levels of total ceramides in MCF-7_ RICTOR^SH^ cells (Figure 2H). Similarly, we also observed >1.2-fold decrease in levels of glucosylceramides and 1.2-1.5 fold increase in levels of ceramides in BT-474_RICTOR^SH^ cells as compared to BT-474 cells (**Supplementary Figure S2G**). As UGCG is responsible for synthesis of glucosylceramides from ceramides, we observed a 2-fold (*p* < 0.05) decrease in UGCG expression by qRT-PCR in MCF-7_ RICTOR^SH^ cells as compared to MCF-7 cells (Figure 2I) that was also validated by immunoblotting (Figure 2J, 2K), and immunofluorescence imaging (Figure 2L). Similarly, we also observed decreased UGCG expression in BT-474_RICTOR^SH^ cells as compared to BT-474 cells by qRT-PCR (**Supplementary Figure S2H**) and immunoblotting along with ∼2-fold decrease in pAKT^S473^ (*p* < 0.0001) (**Supplementary Figure S2I, S2J**). In contrast, we did not observe any change in expression of GBA1 that hydrolyses glucosylceramides to ceramides (Figure 2J, 2K). Therefore, these results suggest that increase in levels of ceramides upon RICTOR silencing is due to transcriptional downregulation of UGCG.

### RICTOR Silencing Inhibits Tumor Progression via UGCG

To validate the RICTOR-mediated regulation of UGCG and its impact on tumor progression, we compared tumor growth kinetics of MCF-7_RICTOR^SH^ and MCF-7_SCRAM^SH^ cells in NOD SCID mice. MCF-7_RICTOR^SH^ tumors showed >3-fold (*p* < 0.001) decrease in kinetics of growth following as compared to MCF-7_SCRAM^SH^ cells (Figure 2M, 2N). Similarly, BT-474_RICTOR^SH^ tumors showed significantly lower tumor progression as compared to BT-474_SCRAM^SH^ tumors (Figure 2O, 2P). These results suggest that RICTOR regulates expression of UGCG at transcriptional level, and increase in glucosylceramides may enhance the cell proliferation and tumor progression (Figure 2Q). We next asked how RICTOR regulates the UGCG expression, and how UGCG-mediated glucosylceramide synthesis enhances tumor progression (Figure 2Q).

**Figure 2.**
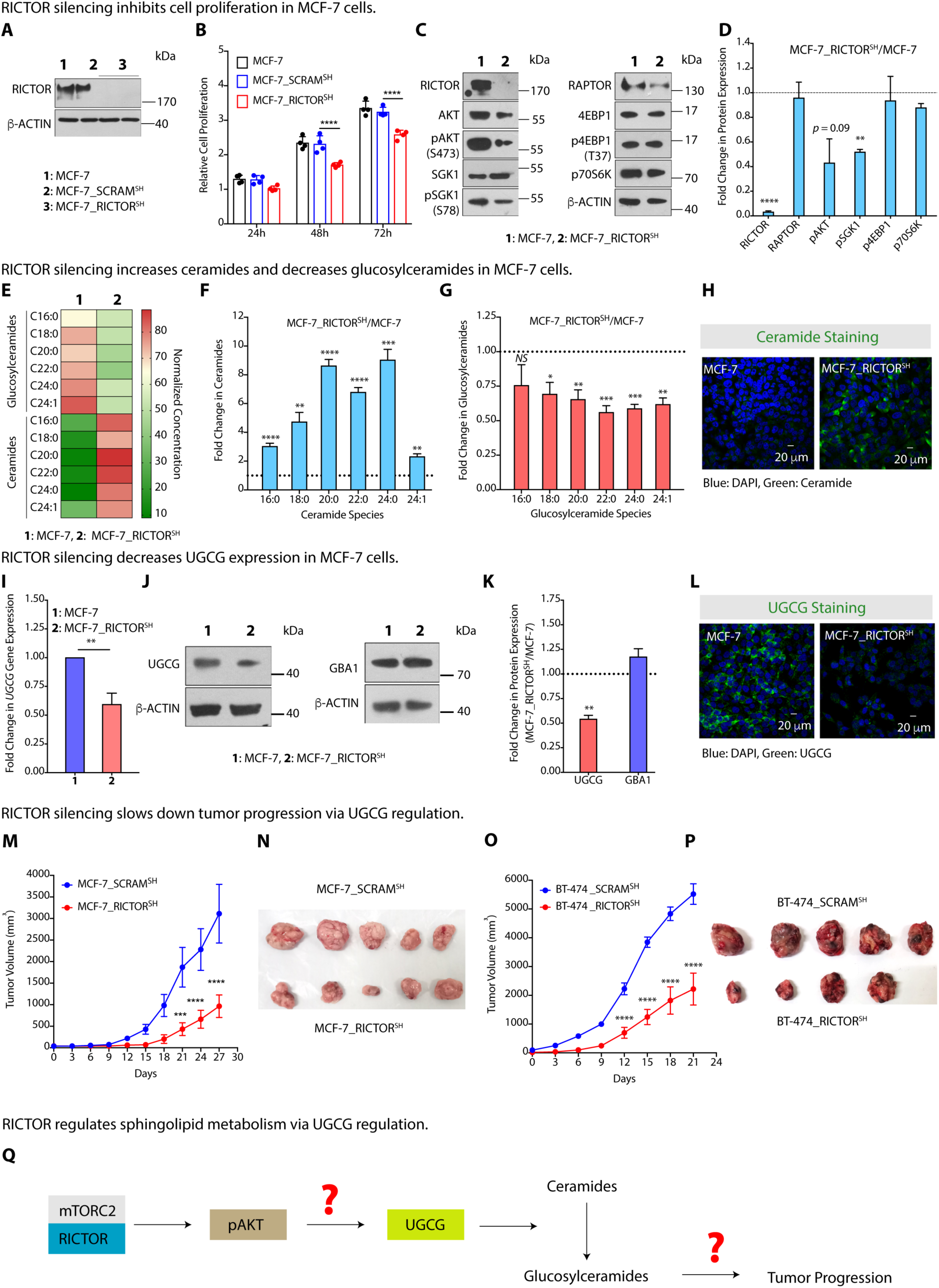
RICTOR Silencing Reduces Glucosylceramide Levels, and Inhibits Tumor Progression. **(A)** Immunoblot with representative β-actin as control confirming knockdown of RICTOR expression in MCF-7_RICTOR^SH^ cells. **(B)** Cell proliferation kinetic studies show decrease in proliferation of MCF-7_RICTOR^SH^ cells (mean ± SD, n = 4) as compared to MCF-7_SCRAM^SH^ cells. **(C, D)** Immunoblots with representative β-actin as control **(C)** and their quantification **(D)** showing changes in protein expression (mean ± SD, n = 3) of RICTOR, RAPTOR, and their downstream effectors. MCF-7_RICTOR^SH^ cells are used as a control. **(E)** Heat map representing normalized absolute quantitation (mean, n = 5) of ceramide and glucosylceramide species (C16:0, C18:0, C20:0, C22:0, C24:0, and C24:1) in MCF-7 and MCF-7_RICTOR^SH^ cells. **(F, G)** Fold change in levels of different sphingolipid species reveal an increase in ceramide levels **(F)** and decrease in glucosylceramide levels **(G)** in MCF-7_RICTOR^SH^ cells as compared to MCF-7 cells. **(H)** Immunofluorescence images confirm enhanced ceramide levels in MCF-7_RICTOR^SH^ cells. **(I, J, K)** qRT-PCR (mean ± SD, n = 5) **(I)**, immunoblots with representative β-actin as control **(J)**, and their quantification (mean ± SD, n = 3) **(K)** demonstrate downregulation of UGCG without any change in GBA1 expression in MCF-7_RICTOR^SH^ cells as compared to MCF-7 cells. **(L)** Immunofluorescence images confirm decreased expression of UGCG in MCF-7_RICTOR^SH^ cells. **(M-P)** Tumor growth kinetics **(M, N)** and pictures of excised tumors **(O, P)** show a significantly slower growth of MCF-7_RICTOR^SH^ (mean ± SEM, n = 5) **(M, O)** and BT-474-7_RICTOR^SH^ (mean ± SEM, n = 5) **(O, P)** tumors as compared to MCF-7_SRAM^SH^ and BT-474_SCRAM^SH^ tumors. **(Q)** A schematic diagram showing the role of putative factors modulating the RICTOR/pAKT-mediated UGCG expression that can lead to altered levels of glucosylceramides, controling tumor progression. Details of all Immunoblots with originals are provided in supplementary information. Data in Figure 2D, 2F, 2G, 2I and 2K were analysed using an unpaired student’s *t* - test, and in Figure 2B, 2M and 2O were analysed using Two-way ANOVA. *p*-value: \**p* < 0.05, \*\**p* < 0.01, \*\*\**p* < 0.001, \*\*\*\**p* < 0.0005.

### RICTOR Regulates UGCG Expression via Transcription Factor Zinc Finger-X linked (ZFX)

We argued that if RICTOR silencing in MCF-7 cells downregulate UGCG expression, then this regulation may be mediated through transcription factors that gets downregulated upon RICTOR silencing. Therefore, we performed differential gene expression analysis on mouse *Rictor* (-) microarray datasets from NCBI GEO, and identified the transcription factors that are downregulated on RICTOR knockdown (Figure 3A) (31). Using bioinformatic analysis, we also identified experimentally validated transcription factors that bind to UGCG promoter in MCF-7 cells (Figure 3A) (32). We found three common transcription factors, Zinc Finger X-linked (*ZFX*), ETS transcription factor (E-74) Like Factor 1 (*ELF1*), and CCCTC-binding Factor (*CTCF*) from above two data sets that may bind to UGCG promoter, and are transcriptionally downregulated on RICTOR silencing. qRT-PCR results confirmed that MCF-7_ RICTOR^SH^ cells have reduced expression of all these three transcription factors (Figure 3B). We further validated the binding of ZFX on UGCG promoter by ChiP-qPCR, and observed ∼1.5-fold (*p* < 0.05) decrease in recruitment of ZFX on UGCG promoter in MCF-7_RICTOR^SH^ cells as compared to MCF-7 cells (Figure 3C**, Supplementary Figure S3A**). In line with this, MCF-7_RICTOR^SH^ cells showed ∼1.5-fold (*p* < 0.05) decrease in ZFX protein expression as compared to MCF-7 cells (Figure 3D, 3E). Similarly, we observed ∼2-fold decrease in ZFX recruitment to UGCG in BT-474_RICTOR^SH^ cells as compared to BT-474 cells (**Supplementary Figure S3B**) along with decrease in ZFX protein expression (**Supplementary Figure S3C**).

### ZFX Regulates RICTOR-mediated UGCG expression

ZFX is a highly conserved Zinc figure protein and oncogenic transcription factor residing on the X chromosome, and is overexpressed in many cancers (33–35). To functionally validate the role of ZFX in UGCG regulation, we used ZFX-overexpressed MCF-7 cells (MCF-7_ZFX^OE^) and only vector overexpressed (MCF-7_VECT^OE^) cells as control. We also used siRNA-mediated ZFX-silenced MCF-7 cells (MCF-7_ZFX^SL^) along with scrambled siRNA-transfected cells (MCF-7_SCRAM^SL^) as control. qRT-PCR results confirmed overexpression of ZFX in MCF-7_ZFX^OE^ cells, and downregulation of ZFX in MCF-7_ZFX^SL^ cells (**Supplementary Figure S3D**) that was further validated by immunoblot (Figure 3F, 3G). Quantitative estimation of ceramides and glucosylceramides by LC-MS/MS showed 1.4-1.8 fold decrease in ceramide levels, and 2-8 fold increase in glucosylceramide levels in MCF-7_ZFX^OE^ cells as compared to MCF-7_VECT^OE^ cells (Figure 3H, 3I). In contrast, we observed a 1.8-3.8-fold increase in ceramide levels, and a 1.5-3.0-fold decrease in glucosylceramide levels on ZFX silencing (Figure 3H, 3I) that was further validated by ceramide staining (Figure 3J). These alterations in ceramide and glucosylceramide levels on ZFX overexpression and silencing prompted us to quantify the UGCG expression. As expected, MCF-7_ZFX^OE^ cells showed ∼2-fold (*p* < 0.05) increase, and MCF-7_ZFX^SL^ cells showed ∼2-fold (*p* < 0.05) downregulation of UGCG expression by qRT-PCR (**Supplementary Figure S3D**) that was validated by immunoblot (Figure 3K, 3L) and immunofluorescence imaging for UGCG (Figure 3M). Similarly, we also observed that siRNA-mediated ZFX silencing in BT-474 cells downregulates UGCG expression (**Supplementary Figure S3E, S3F**), and overexpression of ZFX enhances the UGCG expression (**Supplementary Figure S3G, S3H**).

### ZFX-mediated UGCG Regulation Controls Tumor Progression

To elucidate the effect of ZFX-mediated UGCG regulation on cell proliferation and tumor progression, we compared cell proliferation rates of MCF-7_ZFX^OE^ and MCF-7_ZFX^SL^ cells. MCF-7_ZFX^OE^ exhibited a significant increase in cell proliferation as compared to MCF-7_VECT^OE^ cells, and ZFX silencing showed >1.4-fold (*p* < 0.001) decrease in proliferation as compared to MCF-7_SCRAM^SL^ cells after 72h (**Supplementary Figure S3I**). Scratch wound assay showed >30% increase (*p* < 0.0001) in cell migration in MCF-7_ZFX^OE^ cells, whereas ZFX silencing marked a ∼4-fold decrease (*p* < 0.0001) after 36h (**Supplementary Figure S3J**) that was further validated by transwell migration assay (**Supplementary Figure S3K**). Similarly, we also observed a significant decrease in cell proliferation on ZFX silencing in BT-474 cells (**Supplementary Figure S3L**), and overexpression of ZFX enhanced the cell proliferation (**Supplementary Figure S3M**). Animal studies recorded a significantly higher growth rate of MCF-7_ZFX^OE^ tumors as compared to MCF-7_VECT^OE^ tumors (Figure 3N) that was confirmed by Ki67 staining (Figure 3O).

To validate UGCG-mediated enhanced cell proliferation and tumor progression upon ZFX overexpression, we determined cell proliferation rates of MCF-7_ZFX^OE^ cells upon siRNA-mediated UGCG silencing. Proliferation kinetics demonstrated that UGCG silencing causes a decrease in proliferation rate of MCF-7_ZFX^OE^ cells (Figure 3P). Similarly, nanogel-mediated delivery of UGCG siRNA caused >1.4-fold (*p* < 0.0001) decrease in tumor growth kinetics of MCF-7_ZFX^OE^ tumors (Figure 3Q). Similarly, siRNA-mediated UGCG silencing inhibited the cell proliferation of BT-474-ZFX^OE^ cells (**Supplementary Figure S3N**). Therefore, these results affirm that ZFX, regulated by RICTOR, is one of the transcription factors that modulate UGCG expression in luminal representative MCF-7 and BT-474 cells, priming the increase in glucosylceramide levels, and enhancing tumor progression (Figure 3R). Therefore, the next step was to delineate how RICTOR controls ZFX-mediated UGCG expression (Figure 3R). To see if ZFX-mediated UGCG regulation is a general phenomenon, we silenced ZFX in HCT-116, HepG2, and MDA-MB-453 cancer cells, and observed significant downregulation of UGCG expression (**Supplementary Figure S3O-Q**).

**Figure 3.**
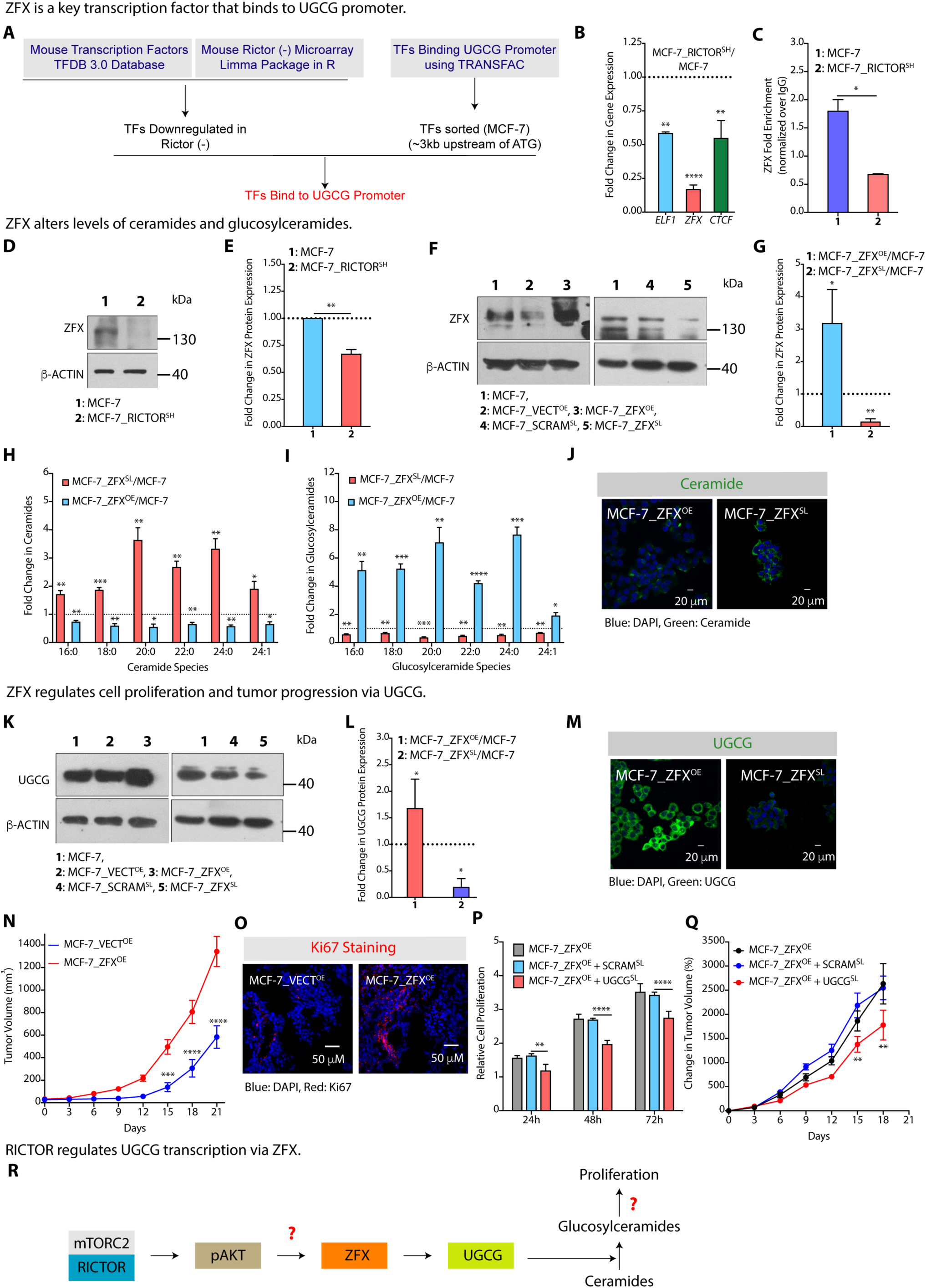
RICTOR Regulates UGCG Expression via Transcription Factor Zinc Finger-X linked (ZFX). **(A)** A schematic diagram showing the workflow used for identification of RICTOR-regulated transcription factors that can bind to UGCG promoter. **(B)** qRT-PCR confirm reduced expression of RICTOR-regulated *ELF1*, *ZFX*, and *CTCF* transcription factors in MCF-7_RICTOR^SH^ cells. **(C)** ChIP-qPCR (mean ± SD, n = 3) results show reduced binding of ZFX to UGCG promoter in MCF-7_RICTOR^SH^ cells. **(D, E)** Immunoblots with representative β-actin as control **(D)** and their quantification (mean ± SD, n = 3) **(E)** show downregulation of ZFX expression in MCF-7_RICTOR^SH^ cells. **(F, G)** Immunoblots with representative β-actin as control **(F)** and their quantification (mean ± SD, n = 3) **(G)** confirm overexpression and silencing of ZFX in MCF-7_ZFX^OE^ and MCF-7_ZFX^SL^ cells respectively. **(H, I)** Fold change (mean ± SEM, n = 5) in absolute levels of ceramides **(H)** and glucosylceramides **(I)** confirm decrease in ceramide levels and increase in glucosylceramide levels in MCF-7_ZFX^OE^ cells. In contrast, MCF-7_ZFX^SL^ cells show higher ceramide levels and reduced glucosylceramide levels. **(J)** Immunofluorescence staining support low ceramide levels in MCF-7_ZFX^OE^ cells, and high ceramide levels in MCF-7_ZFX^SL^ cells. **(K, L)** Immunoblots with representative β-actin as control **(K)** and their quantification (mean ± SD, n = 3) (**L**) witness upregulation and downregulation of UGCG upon overexpression and silencing of ZFX in MCF-7_ZFX^OE^ and MCF-7_ZFX^SL^ cells respectively. **(M)** Immunofluorescence staining support high UGCG expression in MCF-7_ZFX^OE^ cells, and lower expression in MCF-7_ZFX^SL^ cells. **(N)** Tumor growth kinetics recorded a significantly higher growth of MCF-7_ZFX^OE^ (mean ± SD, n = 4) tumors as compared to MCF-7_VECT^OE^ tumors. **(O)** Immunofluorescence images reveal increased expression of Ki67 in MCF-7_ZFX^OE^ tumor sections as compared to MCF-7_VECT^OE^ tumor sections. **(P, Q)** siRNA-mediated silencing of UGCG leads to significant decrease in proliferation of MCF-7_ZFX^OE^ cells **(P)** and reduced tumor growth kinetics in MCF-7_ZFX^OE^ tumors **(Q)**. **(R)** A schematic diagram showing the role of different mechanisms modulating the RICTOR/pAKT-mediated regulation of ZFX/UGCG that leads to altered glucosylceramide levels and controls tumor progression. Details of all Immunoblots with originals are provided in supplementary information. Data in Figure 3B, 3C, 3E, 3G-I, and 3L were analysed using an unpaired student’s *t* -test, and in Figure 3N, 3P, and 3Q were analysed using Two-way ANOVA. *p*-value: \**p* < 0.05, \*\**p* < 0.01, \*\*\**p* < 0.001, \*\*\*\**p* < 0.0005.

### RICTOR Regulates UGCG Expression via Epigenomic Alterations

pAKT-mediated regulation of DNA methyltransferases (DNMTs), histone demethylases (HDMs), and histone methyltransferases (HMTs) can lead to epigenomic alterations, and influence the gene transcription (36). UGCG promoter has multiple CpG islands, and ZFX is known to bind at CpG islands in gene promoters (**Supplementary Figure S4A**) (37). Therefore, we hypothesized that pAKT-mediated regulation of DNMTs may interfere with binding of ZFX to UGCG promoter (38) (Figure 4A). We, first, checked the DNMT expression in MCF-7_RICTOR^SH^ cells, and found high expression of DNMT1, DNMT3A, and DNMT3B in comparison to MCF-7 cells (Figure 4B). Similarly, we also found enhanced expression of DNMT1, DNMT3A, and DNMT3B in BT-474_RICTOR^SH^ cells in comparison to BT-474 cells (**Supplementary Figure S4B**). This enhanced expression may be responsible for increased DNA methylation, and reduced UGCG expression in MCF-7_RICTOR^SH^ cells.

To validate DNMT-mediated UGCG regulation, we inhibited DNMTs by decitabine (DAC) treatment (2, 5, 10 μM), and found 2-4-fold (*p* < 0.05) increase in *UGCG* expression by qRT-PCR (Figure 4C). Immunoblot studies confirmed increase in UGCG expression in MCF-7_ RICTOR^SH^ cells on DAC treatment (Figure 4D). Similarly, we observed an increase in UGCG expression in BT-474_RICTOR^SH^ cells on DAC treatment (**Supplementary Figure S4C**). Using ChIP-qPCR, we witnessed a 2-fold (*p* < 0.05) increase in ZFX enrichment and binding to *UGCG* promoter in DAC-treated MCF-7_RICTOR^SH^ cells (Figure 4E). Therefore, DAC (DNMT inhibition)-mediated upregulation of UGCG is due to enhanced recruitment of ZFX at UGCG promoter site. We further confirmed increase in UGCG expression (Figure 4F) and decrease in ceramides (Figure 4G) by immunofluorescence staining in MCF-7_RICTOR^SH^ cells on DAC treatment. DAC treated MCF-7_RICTOR^SH^ cells also showed increased cell proliferation (Figure 4H) and cell migration as compared to MCF-7 cells (Figure 4I). Similarly, we also observed enhanced cell proliferation of BT-474_RICTOR^SH^ cells on DAC treatment (**Supplementary Figure S4D**). Therefore, these studies demonstrate that DAC-mediated downregulation of DNMTs in MCF-7_RICTOR^SH^ cells enhanced recruitment of ZFX, and elevated UGCG expression, due to reduced DNMT-mediated methylation of UGCG CpG islands.

### RICTOR Regulates UGCG Expression via Histone Demethylase KDM5A

Promoter-associated trimethylation of histone H3 (H3K4Me3) is one of the key targets of PI3K/AKT that acts as a mode of regulating transcriptional competence (39). H3K4Me3 status is regulated by histone demethylase, KDM5A (40). AKT-mediated phosphorylation of KDM5A is instrumental in its nuclear exit leading to elevated H3K4Me3 levels, thereby augmenting gene transcription (Figure 4J). Inhibition of KDM5A using a chemical inhibitor, KDOAM-25, demonstrated an increase in *UGCG* gene expression by 1.5-2.0 fold (p < 0.05) (Figure 4K). This was validated by immunoblot studies showing an increase in UGCG expression with concurrent decrease in KDM5A expression (Figure 4L). ChIP-PCR results confirmed that MCF-7_RICTOR^SH^ cells with reduced AKT activation possess a reduced H3K4Me3 mark as compared to MCF-7 cells, and it was reverted upon KDM5A inhibition (30 μM) with a >1.3-fold (*p* = 0.057) increase (Figure 4M). Immunofluorescence imaging further confirmed increase in UGCG expression (Figure 4N), and decrease in levels of ceramides upon KDM5A inhibition (30 μM) (**Supplementary Figure S4E**).

As KDM5A inhibition causes an increase in UGCG expression and decreases ceramides, cellular assays confirmed increased cell proliferation (Figure 4O) and cell migration in MCF-7_ RICTOR^SH^ cells (Figure 4P) on KDM5A inhibition. Similarly, KDM5A inhibition also causes an increase in cell proliferation of BT-474_RICTOR^SH^ cells (**Supplementary Figure S4F**). We further validated the effect of KDM5A inhibition on UGCG regulation and tumor progression in NOD SCID mice where KDM5A inhibition enhanced tumor growth kinetics of MCF-7_ RICTOR^SH^ cells (Figure 4Q). Therefore, these results confirm that mTORC2-AKT mediated phosphorylation of KDM5A does not allow demethylation of H3K4Me3, and enhanced H3K4Me3 activates UGCG transcription, leading to tumor progression. To complete the circuit connecting the metabolic-gene regulatory signalling, next step was to find how UGCG-mediated increase in glucosylceramides is responsible for enhanced tumor progression (Figure 4R).

**Figure 4.**
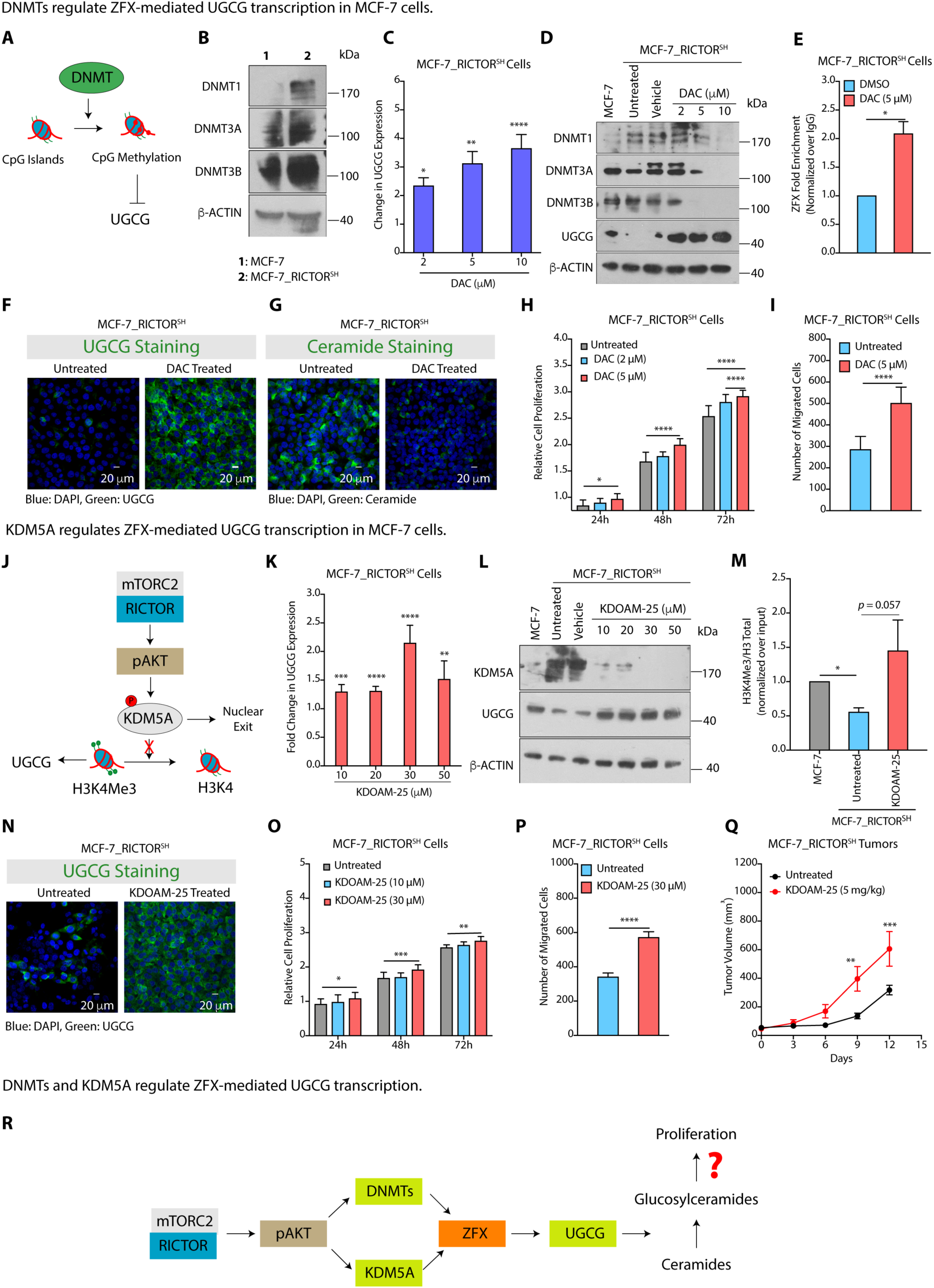
AKT Regulates UGCG Expression via Epigenomic Alterations. **(A)** A schematic diagram showing DNMT-mediated methylation of CpG islands that further regulates transcriptional competence of UGCG. **(B)** Immunoblots with representative β-actin as control showing an increase in expression of DNMTs in MCF-7_RICTOR^SH^ cells as compared to MCF-7 cells. **(C)** qRT-PCR results show an increase in expression of UGCG on inhibition of DNMTs by DAC inhibitor. **(D)** Immunoblots with representative β-actin as control reveal a decrease in expression of DNMTs with a concurrent increase in UGCG expression in MCF-7_RICTOR^SH^ cells upon DAC treatment. **(E)** ChIP-qPCR results (mean ± SD, n = 3) confirm enhanced binding of ZFX to UGCG promoter in MCF-7_RICTOR^SH^ cells on DAC (5 μM) treatment. **(F, G)** Immunofluorescence images confirm enhanced expression of UGCG (**F**) and decreased levels of ceramides **(G)** in MCF-7_RICTOR^SH^ cells on DAC (5 μM) treatment. **(H, I)** Cell proliferation (mean ± SD, n = 3) **(H)** and cell migration (mean ± SD, n = 3) **(I)** assays confirm increased proliferation rate and migration of MCF-7_RICTOR^SH^ cells on DAC (5 μM) treatment. **(J)** A schematic representation of pAKT-mediated regulation of histone demethylase KDM5A that regulates UGCG transcription via histone methylation (H3K4Me3). **(K)** qRT-PCR (mean ± SD, n = 6) results demonstrate an increase in *UGCG* expression in MCF-7_RICTOR^SH^ cells upon treatment with KDM5A inhibitor, KDOAM-25. **(L)** Immunoblot with representative β-actin as control showing an increase in UGCG expression and decrease in KDM5A expression after treatment of MCF-7_RICTOR^SH^ cells with KDOAM-25 inhibitor. **(M)** ChIP-qPCR results (mean ± SD, n = 3) show reduced H3K4me3 mark on UGCG promoter in MCF-7_RICTOR^SH^ cells that increases on treatment with KDOAM-25 inhibitor (30 μM). **(N)** Immunofluorescence images confirm enhanced expression of UGCG in MCF-7_RICTOR^SH^ cells on KDM5A inhibition (30 μM). **(O, P)** Cell proliferation (mean ± SD, n = 3) **(O)** and cell migration (mean ± SD, n = 3) **(P)** assays confirm increased proliferation and migration of MCF-7_RICTOR^SH^ cells on KDM5A inhibition. **(Q)** Inhibition of KDM5A causes a significant increase in tumor growth kinetics of MCF-7_RICTOR^SH^ cells. **(R)** A schematic diagram showing pAKT substrates DNMTs and KDM5A that epigenetically regulate ZFX enrichment on UGCG promoter, and regulate UGCG transcription leading to altered levels of glucosylceramides and cell proliferation. Details of all Immunoblots with originals are provided in supplementary information. Data in Figure 4C, 4E, 4I, 4K, 4M, and 4P were determined using an unpaired student’s *t* -test, and in Figure 4H, 4O, and 4Q were analysed using Two-way ANOVA. *p*-value: \**p* < 0.05, \*\**p* < 0.01, \*\*\**p* < 0.001, \*\*\*\**p* < 0.0005.

### ZFX-mediated UGCG Regulation Alters the Level of Gangliosides

Synthesis of glucosylceramides is followed by synthesis of lactosylceramides and monosialic acid containing GM3 gangliosides (Figure 5A). GM3 gangliosides can either be converted to monosialic acid gangliosides, GM2 and GM1, or disialic acid ganglioside GD3 (Figure 5A). As RICTOR controls ZFX-mediated transcriptional regulation of UGCG and synthesis of glucosylceramides, we quantified the impact of ZFX-UGCG regulation on levels of lactosylceramides and three key gangliosides, GM3, GD3, and GM1 in MCF-7, MCF-7_RICTOR^SH^, MCF-7_UGCG^OE^, and MCF-ZFX^OE^ cells. As RICTOR silencing reduces the levels of glucosylceramides, we observed >4-fold (*p* < 0.005) decrease in C16:0 lactosylceramides in MCF-7_RICTOR^SH^ cells as compared to MCF-7 cells (Figure 5B). In contrast, overexpression of ZFX and UGCG induced a 4-8 fold increase in lactosylceramides (Figure 5B).

MCF-7_RICTOR^SH^ cells have significantly reduced levels of all species of GM3 gangliosides (C16:0, C18:0, C20:0, C22:0, and C24:0) that may be due to reduced levels of glucosylceramides and lactosylceramides (**Supplementary Figure S5A,** Figure 5C). In contrast, there was a several-fold increase in levels of all species of GM3 gangliosides in MCF-7_UGCG^OE^ and MCF-ZFX^OE^ cells due to higher expression of glucosylceramides (**Supplementary Figure S5A**). As C16:0 GM3 gangliosides are most abundant in MCF-7 cells, and it showed a sharp increase in MCF-7_UGCG^OE^ and MCF-7_ZFX^OE^ cells (Figure 5C), we compared the changes in levels of C16:0-derived GM1 and GD3 gangliosides among different cell types. MCF-7_RICTOR^SH^ cells showed >3-fold (*p* < 0.05) decrease in levels of GM1 gangliosides (Figure 5D), and ∼ 2.5 - fold (*p* < 0.05) decrease in levels of GD3 gangliosides as compared to MCF-7 cells (Figure 5E). This reduction in GM1 and GD3 gangliosides on RICTOR silencing is due to reduced levels of glucosylceramides and GM3 gangliosides. In contrast, there was 4-10-fold decrease in GM1 gangliosides (Figure 5D), and 1.5-2.0 fold increase in GD3 gangliosides in MCF-7_UGCG^OE^ and MCF-7_ZFX^OE^ cells over MCF-7 cells (Figure 5E). Interestingly, MCF-7_RICTOR^SH^ cells have >2-fold higher levels of GM1 gangliosides, and >6-fold lower levels of GD3 gangliosides than MCF-7_UGCG^OE^ and MCF-7_ZFX^OE^ cells (Figure 5D, 5E). These alterations in GM1 and GD3 gangliosides were then validated by flow cytometry (Figure 5F, 5G). Similarly, flow cytometry analysis showed lower expression of GM1 gangliosides and enhanced expression of GD3 gangliosides in BT-474_UGCG^OE^ and BT-474_ZFX^OE^ cells in comparison to BT-474 cells, whereas BT-474_RICTOR^SH^ cells have higher expression of GM1 gangliosides and lower GD3 levels (**Supplementary Figure S5B, 5C**). To reaffirm that UGCG is responsible for these changes in level of gangliosides, we evaluated the expression of other ganglioside-metabolic pathway enzymes (ST8SIA-V, B4GALNT1, B3GALT4, ST8SIA1), and observed no appreciable change in MCF-ZFX^OE^ cells over MCF-7 cells (**Supplementary Figure S5D**).

### GD3 Gangliosides cause EGFR-mediated Tumor Progression

Gangliosides present in GEMs are well-known to regulate RTK signalling that is contingent upon nature and relative quantity of gangliosides, kind of growth factor receptors, and cell type (41). Gangliosides like GD2 and GD3 can activate RTK signalling, and enhance cancer cell proliferation (42). In contrast, gangliosides like GM1 can mitigate RTK signalling via inhibiting the dimerization of growth factor receptors (43). To study the regulatory effects of alterations in gangliosides on RTK signalling, we estimated the levels of phosphorylated epidermal growth factor receptor (EGFR) and its downstream signalling components. Immunoblot analysis showed no alterations in levels of total EGFR in MCF-7_RICTOR^SH^, MCF-7_UGCG^OE^, and MCF-ZFX^OE^ cells as compared to MCF-7 cells (Figure 5H). However, we observed a >3-fold (*p* < 0.05) increase in phosphorylated EGFR (EGFR^Y1173^ and EGFR^Y1068^) in MCF-7_UGCG^OE^ and MCF-7_ZFX^OE^ cells having higher levels of GD3 gangliosides over that of MCF-7 cells (Figure 5H, 5I). Similarly, we observed activation and upregulation of downstream signalling intermediates, pAKT^S473^, and extracellular signal-regulated protein kinase/p(ERK1/2) in MCF-7_UGCG^OE^ and MCF-7_ZFX^OE^ cells in comparison to MCF-7 cells (Figure 5H, 5I). Therefore, these results suggest that overexpression of ZFX or UGCG causes increased activation of EGFR-mediated RTK signalling. Similarly, we observed the activation of EGFR signalling in BT-474_UGCG^OE^ and BT-474_ZFX^OE^ cells in comparison to BT-474 cells (**Supplementary Figure S5E**).

To validate GD3 ganglioside-mediated activation of EGFR, we performed siRNA-mediated silencing of GD3 synthase (ST8SIA1) in MCF-7_ZFX^OE^ cells, and observed a significant downregulation in pEGFR^Y1173^ and pEGFR^Y1068^ levels, though total EGFR level was unaltered (Figure 5J, 5K). ST8SIA1 silencing using siRNA also attenuated the cell proliferation of MCF-7_ZFX^OE^ cells (Figure 5L), and even abrogated the tumor growth kinetics in mice xenografts (Figure 5M). Similarly, ST8SIA1 silencing using siRNA also attenuated the cell proliferation of BT-474_ZFX^OE^ cells (**Supplementary Figure S5F**). To validate GD3 ganglioside mediated activation and proliferation of cancer cells, we incubated MCF-7_RICTOR^SH^ cells with GD3 gangliosides, and observed >1.5 fold increase in cell proliferation (Figure 5N), and incubation of MCF-7_UGCG^OE^ and MCF-ZFX^OE^ cells with GM1 gangliosides inhibited the cell proliferation (Figure 5O). Similarly, incubation of BT-474_RICTOR^SH^ cells with GD3 gangliosides enhanced the cell proliferation (**Supplementary Figure S5G**), and incubation of BT-474_UGCG^OE^ and BT-474_ZFX^OE^ cells with GM1 gangliosides inhibited the proliferation (**Supplementary Figure S5H**). Therefore, these results confirm that GD3-mediated activation of EGFR is responsible for cell proliferation and tumor progression, thus completing the metabolic-signalling-gene regulation circuit. (Figure 5P).

**Figure 5.**
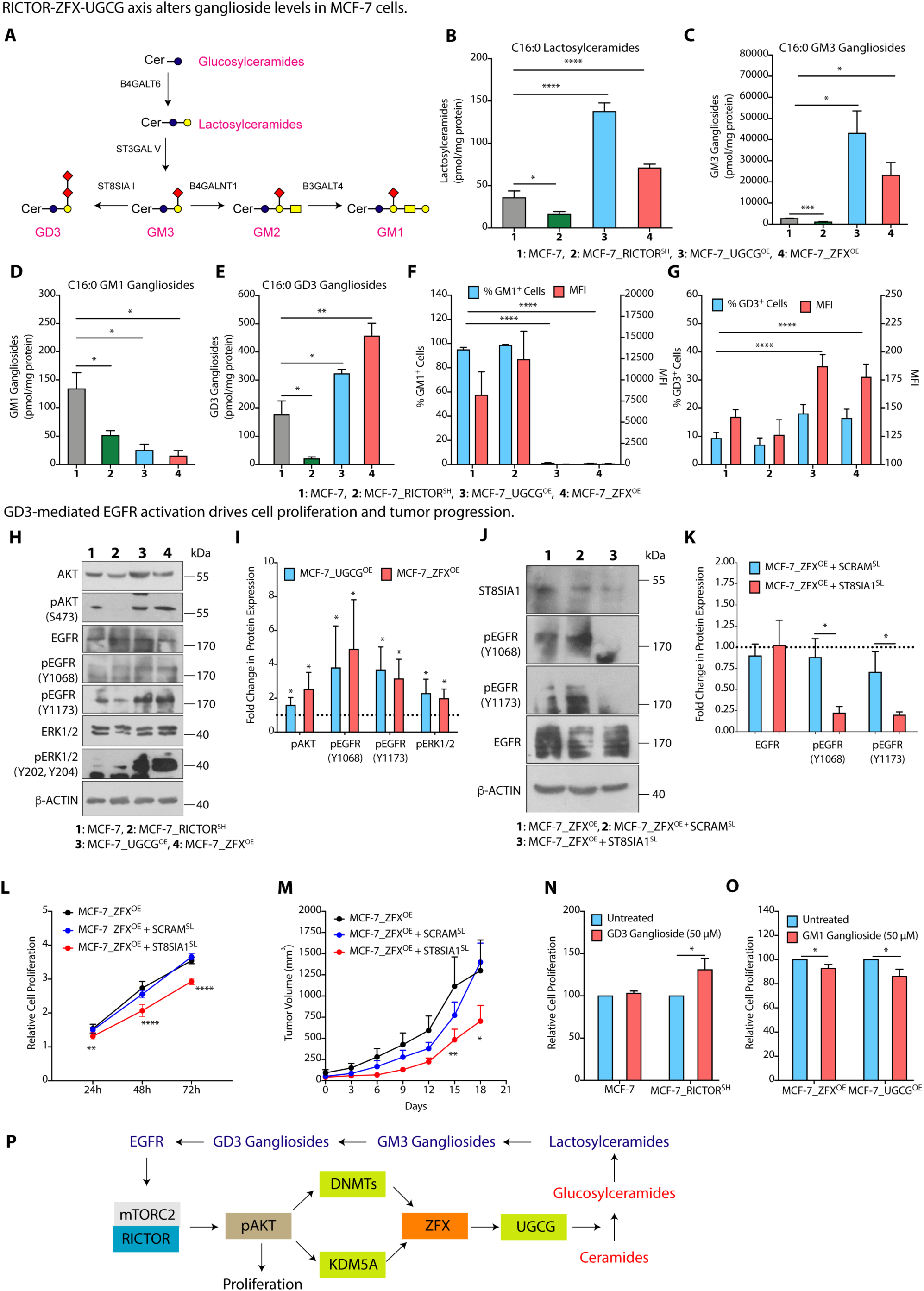
ZFX-mediated UGCG Regulation Alters Ganglioside Levels and Activates EGFR. **(A)** Schematic presentation of a part of ganglioside metabolic pathway showing synthesis of GD3 and GM2/GM1 gangliosides from common precursor GM3 gangliosides. **(B-E)** Absolute quantitation (mean ± SEM, n = 5) of C16:0-derived lactosylceramides **(B)**, GM3 **(C)**, GM1 **(D)**, and GD3 **(E)** gangliosides show increase in GM3 and GD3 ganglioside levels, and decrease in GM1 ganglioside levels in MCF-7_UGCG^OE^ and MCF-7_ZFX^OE^ cells. **(F, G)** Quantification of GM1^+^ **(F)** and GD3^+^ **(G)** cells and MFI (mean fluorescence intensity) by flow cytometry confirm decrease in GM1 expression and enhanced GD3 expression in MCF-7_UGCG^OE^ and MCF-7_ZFX^OE^ cells. **(H, I)** Immunoblots along with representative β-actin as control **(H)** and their quantification (mean ± SD, n = 3) **(I)** reveal an increase in levels of pEGFR^Y1173^, pEGFR^Y1068^, pAKT^S473^, and pERK1/2^(Y202,Y204)^ in MCF-7_UGCG^OE^ and MCF-7_ZFX^OE^ cells as compared to MCF-7 cells. **(J, K)** Immunoblots with representative β-actin as control **(J)** and their quantification (mean ± SD, n = 3) (**K**) show attenuated EGFR activation on siRNA-mediated silencing of GD3 synthase (ST8SIA1). **(L)** Cell proliferation assay demonstrates decrease in cell proliferation (mean ± SD, n = 3) of MCF-7_ZFX^OE^ cells on siRNA-mediated inhibition of ST8SIA1. **(M)** Tumor growth kinetics using xenograft studies show a decrease in growth kinetics (mean ± SEM, n = 5-6) of MCF-7_ZFX^OE^ tumors on siRNA-mediated inhibition of GD3 synthase (ST8SIA1). **(N, O)** Cell proliferation assay showing increase in proliferation of MCF-7_RICTOR^SH^ cells upon feeding with GD3 gangliosides **(N)**, and decrease in cell proliferation upon feeding of MCF-7_UGCG^OE^ and MCF-7_ZFX^OE^ with GM1 gangliosides **(O)**. **(P)** A schematic diagram showing the role of ZFX-mediated transcriptional regulation of UGCG that modulates ganglioside biosynthesis and ganglioside-mediated EGFR activation. Details of all Immunoblots with originals are provided in supplementary information. Data in Figure 5G, 5I, 5K, 5N, and 5O) were analysed using an unpaired student’s *t* -test, and in Figure 5L and 5M was analysed using Two-way ANOVA. *p*-value: \**p* < 0.05, \*\**p* < 0.01, \*\*\**p* < 0.001, \*\*\*\**p* < 0.0005.

### ZFX Expression is Strongly Associated with UGCG in Luminal Patients

To find the association between ZFX and UGCG in luminal breast cancer patients, we analyzed the TCGA-BRCA and METABRIC patient datasets. TCGA patient dataset (N = 1082) based on PAM50 classification (N = 833) was divided into luminal A (N = 416), luminal B (N = 185), and non-luminal subtypes (N = 232) (Figure 6A) (44). Differential gene expression data analysis showed that luminal A and B tumor tissues have significantly higher expression of UGCG (Figure 6B) and ZFX (Figure 6C) as compared to Basal and HER2^+^ groups. Correlating ZFX and UGCG expression to ER and PR status revealed that ER^+^ (Figure 6D, 6E) and PR^+^ (Figure 6F, 6G) tumors have significantly higher expression of UGCG (Figure 6D, 6F) and ZFX (Figure 6E, 6G). Gene expression analysis further revealed that >56% of luminal (luminal A and B) patients have high expression of both UGCG and ZFX (Figure 6H). Similarly, we divided the METABRIC data sets based on PAM50 classification (N = 1905) into Luminal A (N = 696), Luminal B (N = 474), and non-Luminal (N = 728) subtypes (**Supplementary Figure S6A**) (45). Metadata analysis further confirmed that luminal A and luminal B tumor tissues have higher UGCG and ZFX expression over other subtypes as observed in TCGA data set (**Supplementary Figure S6B, S6C),** and expression of UGCG and ZFX is also high in ER^+^ and PR^+^ tumor tissues (**Supplementary Figure S6D-G).** We identified 45% of luminal (luminal A and B) patients with higher UGCG expression also exhibit higher ZFX expression (**Supplementary Figure S6H**).

To further validate the association of ZFX and UGCG in breast cancer patients, we quantified the expression of UGCG and ZFX by immunohistochemical (IHC) analysis in tumor samples of all subtypes (N = 90) (**Dataset 2**). In concurrence with TCGA BRCA and METABRIC datasets, IHC analysis confirmed that ∼15% of luminal patients have high ZFX and UGCG expression (Figure 6I, J). Finally, we quantified the expression of *UGCG* and *ZFX* from luminal tumors by qRT-PCR, and observed ∼2-fold (*p* < 0.05) increase in expression of ZFX and UGCG in tumor tissues over adjacent matched normal control (Figure 6K). Therefore, these results support a positive correlation between ZFX and UGCG expression in luminal patients.

**Figure 6.**
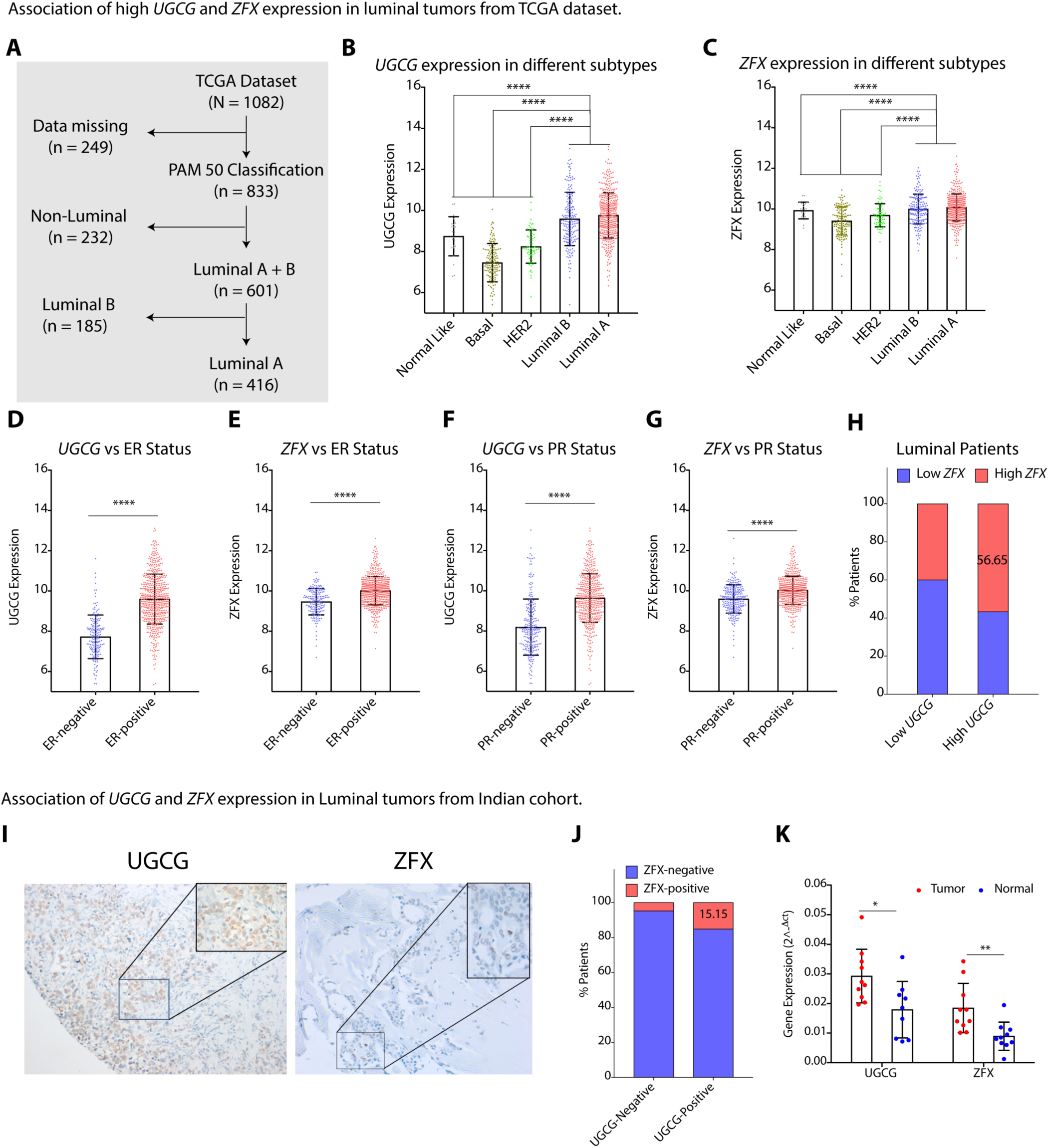
ZFX Expression is Strongly Associated with UGCG in Luminal Patients. **(A)** A schematic diagram showing the PAM50 classification of TCGA tumor dataset used for analysis. **(B, C)** Gene expression of *UGCG* **(B)** and *ZFX* **(C)** in different breast cancer subtypes (PAM50) of TCGA dataset confirms high expression of *UGCG* and *ZFX* in luminal subtypes as compared to other subtypes. (**D-G**) Change in expression of *UGCG* **(D, F)** and *ZFX* **(E, G)** with respect to ER **(D, E)** and PR **(F, G)** status in breast tumors of TCGA dataset confirm high *UGCG* **(D, F)** and high *ZFX* **(E, G)** expression in ER^+^ and PR^+^ tumors. **(H)** Percentage of tumors having high expression of *UGCG* and *ZFX* among luminal subtype tumors in TCGA data set. **(I)** Representative immunohistochemical images show enhanced cytoplasmic UGCG and increased nuclear stain of ZFX in luminal breast tissues. All images are at 100X magnification, and insets are at 400x magnification. (**J**) Percentage of luminal tumors from Indian cohort (N = 90) positive for both UGCG and ZFX on immunohistochemical staining. **(K)** qRT-PCR (mean ± SD, n = 10) validation showing high *UGCG* and high *ZFX* expression from luminal subtype tumors in comparison to adjacent normal tissues in an Indian cohort. Data in Figure 6B and 6C were analyzed using One-way ANOVA, and data in Figure 6D-6G and 6K were analyzed by an unpaired student’s *t* -test. *p*-value: \**p* < 0.05, \*\**p* < 0.01, \*\*\*\**p* < 0.0001.

## DISCUSSION

Treating cancer is like the Herculean duel with the chthonic creature Hydra, whose decapitation magically led to a botanical duplication of the regenerated heads. The myriad of alternative strategies that cancer cells employ to achieve survival advantage over the clinical interventions is a similar saga. One of the prime reason for this is our lack of complete understanding of the metabolic signalling and gene regulatory networks that cancer cells deploy in order to survive. More important is to understand how these networks are interconnected so that multiple nodes in the circuit/network can be combinatorially targeted to overrule their survival strategies. In this context, herein, we have mapped the first step of metabolic-signalling-gene regulatory circuit connecting ganglioside metabolism with cancer, controlled by EGFR-mTORC2/RICTOR complex (Figure 7).

The major biological roles of sphingolipids and gangliosides at the cell surface include modulating the lipid phase of cellular membranes, acting as ligands for membrane receptors, kinases, and enzymes, and surface recognition through glycan interactions by glycosphingolipids (46). Gangliosides, as a part of GEMs, can act as double-edged swords where they can either augment or inhibit the growth factor-mediated cell proliferation through activation/deactivation of RTK signalling cascades (47). Using an elegant precedence of ganglioside-mediated activation of growth factor receptor signalling, our work reveals that increased GD3 and GM3 levels and reduced GM1 levels in response to altered expression of UGCG boost the EGFR autophosphorylation status, and subsequent downstream growth signalling in luminal cancers. Although RTK inhibitors have made breakthroughs in tumor treatment options, however, RTK co-activation networks mark a serious limitation in their use (48). This effectively suggests that systematic effort in manipulation of gangliosides via UGCG or GD3/GM3 synthesizing enzymes can serve as a strategy to prevent activation of multiple RTKs that network for accelerated tumor growth in luminal subtype.

Targeting RICTOR downstream of RTKs may be another node that can be tapped simultaneously. We ruled out the cross-talk of mTORC1 on RICTOR silencing, as RAPTOR and downstream targets like pS6 Kinase and p4EBP1 were unchanged in MCF-7_RICTOR^SH^ cells. However, there are many negative feedback loops working between mTOR cascades which though important, investigating all of these were beyond the scope of this study. Although many studies till now have reported high UGCG expression in ER^+^ luminal tumors (49, 50), but none delved into its molecular mechanism. Our study on the other hand unravelled and validated that, ZFX, a key C2H2-type, ZNF family transcription factor, mediates *UGCG* expression, and thereby modulates sphingolipid and ganglioside metabolism, and luminal tumor progression. Thus, to avoid the activation of mTOR-mediated negative feedback loops promoting cell proliferation, efforts need to be diverted on manipulation of ZFX controlled *UGCG* expression that will mimic the effect of RICTOR inhibition without associated side effects. This regulatory effect of ZFX may be one of the reasons why genetic manipulations to silence ZFX in breast, colorectal, pancreatic and renal cancers reduced proliferation and invasion, and predicted good prognosis (51, 52). It also needs to be emphasized that depending on the cell line and growth signalling pathway involved, there may be multiple transcription factor/s regulating the *UGCG* in a context-dependent manner.

PI3K/AKT signalling stabilizes DNA methyltransferases affecting the global methylation pattern and transcriptional activation of genes in different cancers (53). Transcription factor mediated maintenance of DNMTs to promoters involve co-operative association or recruitment of additional co-repressors (54). In presence of multiple CpG islands in *UGCG* promoter, transcription factor mediated switch of its activator and repressor effect probably shifts the balance of DNMT/ZFX ratio more towards binding of DNMTs (DNMT1, 3A and 3B) on this oncogene promoter that compromises ZFX binding in MCF-7_RICTOR^SH^ cells. However, it remains to be seen which of the DNMTs, if not more than one member, are recruited to *UGCG* promoter that prevents ZFX enrichment when pAKT levels are low. Similarly, AKT-mediated phosphorylation of KDM5A causes it nuclear export, and allows ZFX binding to the promoter by allowing histone methylation. Besides, pAKT-mediated epigenetic regulations can be further exploited to control gene expression mediated metabolic networking. In summary, our study endorses that EGFR-mTORC2-RICTOR-AKT-UGCG-Ganglioside circuit regulates tumor progression in luminal breast cancer cells, and provides us an opportunity to intervene at multiples nodes to tame cancer cells. It is prudent to mention that underlined paradigm of mTORC2/RICTOR regulated expression of *UGCG* impacting the level of glucosylceramides may be part of a fundamental mechanism in breast tumor development especially in luminal tumors as shown in clinical samples, that certainly requires more attention and research.

**Figure 7.**
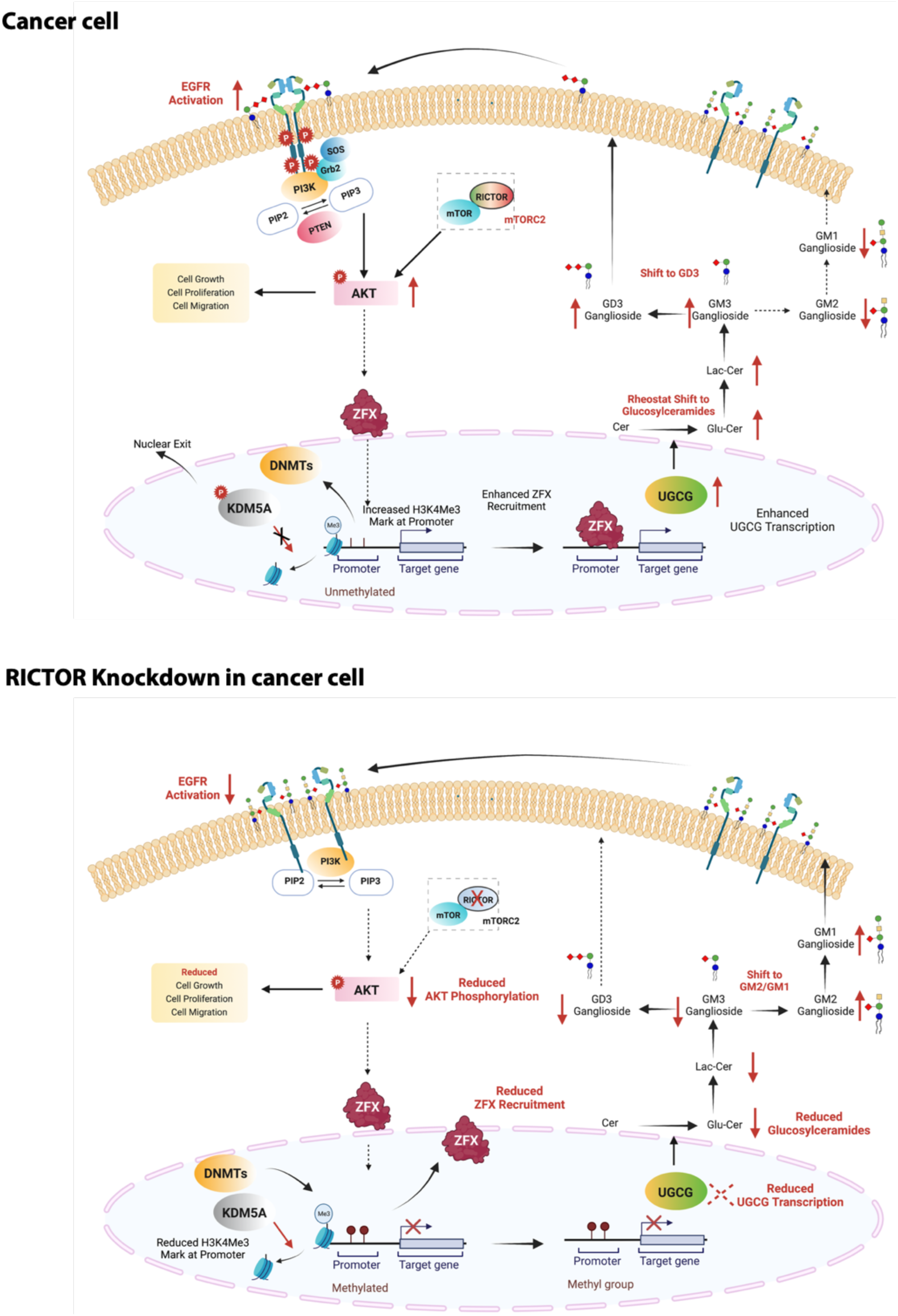
Schematic showing the metabolic-signaling-gene regulatory circuit connecting ganglioside metabolism with cancer. Ganglioside-mediated EGFR activation followed by mTORC2/RICTOR signalling epigenetically modifies UGCG transcription efficiency, ultimately leading to tumor progression.

## MATERIALS and METHODS

### MATERIALS

#### Cell culture

MCF-7 and BT-474 cells (ATCC, USA), DMEM media (Cat# D5648) Sigma, USA, MEBM media (Cat# CC-3151) Lonza, Switzerland, MEM media (Cat# AL081) HiMedia, USA, DPBS (Cat# D5652) Sigma, USA, FBS (Cat# 10270) Gibco, USA, Penicillin-Streptomycin (Cat# 113-98-43810-74-0) HyClone, USA, Lipofectamine 2000, (Cat# 11668019) Invitrogen, USA, Trypsin (Cat# TCL007) HiMedia, USA, Puromycin (Cat# P7255) Sigma, USA, G418 (Cat# A1720) Sigma, USA, Haemocytometer (Cat# Z359629) Bright-Line^TM^, USA, Transwell Migration Plate (Cat#3464) Corning, USA, Crystal violet dye (Cat# C0775) Sigma, USA, shRNA Control (Cat# SHC202V) Sigma, USA, shRNA RICTOR (Cat# SHCLNV) Sigma, USA, UGCG siRNA (Cat# AM51331) Ambion, USA, ZFX siRNA (Cat#L -006572-00-0005) Dharmacon, USA, Scrambled siRNA (Cat# D-001810-10-05) Dharmacon, USA, GD3 Synthase (ST8SIA1) siRNA (Cat# EHU025731-20UG) Merck, USA, KDM5A siRNA (Cat# EHU012051-20UG) Merck, USA, 5-Aza-2′-deoxycytidine (DAC) (Cat#A3656-10MG) Sigma, USA, KDOAM25 Hydrochloride hydrate (Cat# SML2774-5MG) Sigma, USA.

#### Biochemicals and Kits

Qubit RNA AH Assay Kit (Cat# Q32853) Invitrogen, USA, RNAiso Plus (Cat# 9109) DSS Takara, India, RNeasy^®^ Lipid Tissue Mini Kit (Cat# 74804) Qiagen, Germany, Ethanol (Cat# 100983) Merck, USA, MOPS, free acid (Cat# MB0360) Bio basic, Canada, formaldehyde (Cat# MB059) HiMedia, USA, Ethidium bromide (Cat# E8751) Sigma, USA, 100 bp DNA ladder (Cat# BM001-R500) BR Biochem, China, TURBO^TM^ DNase (Cat# AM2238) Invitrogen, USA, iScript^TM^ cDNA synthesis kit (Cat# 1708891) Bio-Rad, USA, Agarose (Cat# A9539) Sigma, USA, iTaq™ universal SYBR®Green supermix (Cat# 1725124) Bio-Rad, USA, Ethylenediaminetetraacetic acid disodium salt dihydrate (Cat# E5134) Sigma, USA,Tris (Cat#MB029) HiMedia, USA, NaCl (Cat#GRM853) HiMedia, USA, MgCl_2_ (Cat# 208337)Sigma, USA, CaCl_2_ (Cat# 449709) Sigma, USA, Triton X-100 (Cat# T8787) Sigma, USA, Sodium deoxycholate (Cat# 1.06504) Millipore, USA, DTT (Cat# DTT-RO) Roche, Switzerland, complete™ Protease Inhibitor Cocktail (Cat# CO-RO) Roche, Switzerland, SUPERase• In™ RNase Inhibitor (Cat# AM2694) Thermo Scientific, USA, Xylene cyanol FF (Cat# X4126) Sigma, USA, Hydrochloric acid (Cat# 29505) Thermo Fisher, USA, Bromophenol blue sodium salt (Cat# B8026) Sigma, USA, Sodium dodecyl sulfate (Cat# L3771) Sigma, USA, Lithium Chloride (Cat# 9650-100G) Sigma, USA, Acrylamide (Cat# AB1032) Bio basic, Canada, Polyoxyethylenesorbitan monolaurate (Tween® 80) (Cat# GRM156) HiMedia, USA, Polyoxyethylenesorbitan monolaurate (Tween® 20) (Cat# P7949) HiMedia, USA, Glycerol (Cat# GRM1027) HiMedia, USA, Ammonium persulfate (Cat# A3678) Sigma, USA, Phenylmethanesulfonyl fluoride (Cat# P7626) Sigma, USA, 1X Protease inhibitor cocktail (Cat# R1329) Fermentas, USA, HindIII (Cat# R0104S) NEB, USA, Xhol (Cat# R0146S) NEB, USA, NEB Buffer 2.1 (Cat# B7202S) NEB, USA, *N*,*N*′-methylene bisacrylamide (Cat# M7279) Sigma, USA, Sodium chloride (Cat# GRM853), HiMedia, USA, Glycine (Cat# G8898), Sigma, USA, Bovine serum albumin fraction-V, HiMedia, USA, nitrocellulose (Cat# GRM105), Merck, USA, PVDF membrane (Cat# IPVH00010), Merck, USA, Chemiluminescent HRP substrate (Cat# WBKLS0500), Merck, USA, Pierce^TM^ BCA protein assay kit (Cat# 23227), Thermo Scientific, USA, MTT (Cat# M5655), Sigma, USA, Hoechst 33258 (Cat# 861405), Sigma, USA, Paraformaldehyde (Cat# 81847), Thomas baker, India, Cryomatrix (Cat# 6769006), Thermo Scientific, USA, Poly-lysine slides (Cat# P0425), Sigma, USA, Goat serum (Cat# RM10701), HiMedia, USA, Allprotect Tissue Reagent (Cat# 76405), Qiagen, Germany, Prolong gold anti-fade reagent (Cat# P36934), Life technologies, USA, Sectioning blade (Cat# 152580), Micron, India, Triton X-100 (Cat# T9284), Sigma, USA, Taurocholic acid sodium (Cat# T4009), Sigma, USA, citric acid (Cat# 251275), Sigma, USA, Disodium hydrogen orthophosphate dihydrate (Cat# 40158), S.D. Fine, India, sodium hydroxide (Cat# 40167), S.D. Fine, India, Kanamycin (Cat# 25389-94-0), GoldBiocom, India, Dynabead Protein A (Cat# 10002D), Thermo, USA, Dynabead Protein G (Cat# 10004D), Thermo, USA, Plasmid Midi Prep (Cat# 12143), Qiagen, Germany, Polybrene transfection reagent (Cat# T1003), Sigma, USA, developer (Cat# 4908216), Carestream, USA, Fixer (Cat# 4908232), Carestream, USA, XBT X-Ray film (Cat# 6568307), Carestream, USA, Immobilon Western Chemiluminescent HRP (Cat# WBKLS0500), Merck Millipore, USA.

#### Chemicals for lipidomics and mass spectrometry studies

Methanol (MS Grade Cat#34966), Honeywell, USA, Chloroform (MS Grade Cat#25669-1L), Honeywell, USA, 2-Propanol (Cat#34965), Honeywell, USA, Formic Acid (MS Grade Cat# 56302-50ML), Fluka, USA, Ammonium formate, (Cat# 14266-25G), Honeywell Fluka, USA, Acetonitrile (Cat#34967) Honeywell, USA, Ammonium acetate (Cat#14267-25G), Honeywell, USA, Ammonium hydroxide (Cat# 16227), Thermo Fisher, USA, Triethyl ammonium bicarbonate buffer Sodium Hydroxide (Cat# T70408), Sigma, USA, Sodium Hydroxide (Cat# 13913) SRL Chem, India, Chymotrypsin (Cat# 11418467001), Merck, USA, Water (MS Grade Cat#39253-4L) Riedel-de haen, Germany. Potassium hydroxide (Cat#84749), Sisco Research, India, Iodoacetamide (Cat# 144-48-9) Sigma, USA, Glacial acetic acid (Cat# 144-48-9) Merck, USA, Digitonin (Cat# D141) Sigma, USA, ACQUITY UPLC BEH Shield RP18 column (Cat#186002854) Water Ltd., ChromXP C18-CL trap column (Cat#5016752) was purchased from Eksigent, ABSciex, USA, nanoViper C18 column (Cat# 164569) Thermo Scientific, USA, Kinetex® C18 column (Cat#00B-4601-AN) Phenomenex®, USA, Ceramide/Sphingoid Internal Standard Mixture II (Cat# LM6005-1EA) Avanti Polar Lipids, USA, C18 Ganglioside GM3-d3 (d18:1/18:0d3) (ammonium salt) (Cat# 24850) Cayman Chemicals, USA, Ganglioside GM3 (Bovine Milk) (Cat# 860058P) Avanti Polar Lipids, USA, Ganglioside GM1 (Bovine Brain) (Cat# 860065P) Avanti Polar Lipids, USA, Ganglioside GD3 (Bovine Milk) (Cat# 860060P) Avanti Polar Lipids, USA, FITC labelled cholera toxin B (Cat#1655) Sigma-Aldrich, USA.

#### Antibodies

LASS1 (Cat#H00010715-A01), Abnova, Taiwan, LASS2 (Cat#H00029956-M01A) Abnova, Taiwan, LASS4 (Cat#H00079603-M01) Abnova, Taiwan, LASS5(Cat# ab73289) Abcam, UK, LASS6 (Cat#H00253782-M01) Abnova, Taiwan, A-SMase (SMPD1 Cat# ab83354) Abcam, UK, N-SMase1 (SMPD2 Cat# ab131330) Abcam, UK, N-SMase2 (SMPD3 Cat#ab199399) Abcam, UK, N-SMase3 (SMPD4 Cat#ab133935) Abcam, UK, Ceramide Glucosyl transferase (Cat#ab124296) Abcam, UK, Ceramide Glucosyl transferase IHC (Cat# ab197369) Abcam, UK, Sms1 (Cat#ab135365) Abcam, UK, Sms2 (Cat#ab237681) Abcam, UK, Gba1 (Cat#ab88300) Abcam, UK, Glb1 (Cat#ab96239) Abcam, UK, B4galt6 (Cat#ab200639) Abcam, UK, eIF4EBP1 (Cat# ab32024) Abcam, UK, P-eIF4EBP1 (Cat# ab75767) Abcam, UK, DNMT1 (Cat# ab188453) Abcam, UK, DNMT3A (Cat# ab2851) Abcam, UK, DNMT3B (Cat# ab2851) Abcam, UK, H3K4ME3 (Cat# ab1791) Abcam, UK, H3 (Cat# ab8580) Abcam, UK, Rictor (Cat# 2140S) Cell Signaling, USA, Raptor (Cat# 2280) Cell Signaling, USA, Akt (Cat# 4685) Cell Signaling, USA, P-Akt (Cat# 4060) Cell Signaling, USA, P-SGK1 (Cat# 5599) Cell Signaling, USA, P70 S6 Kinase (Cat# 5599) Cell Signaling, USA, ZFX (Cat# 5419) Cell Signaling, USA, JARID1A (KDM5A) (Cat# 3876S) Cell Signaling, USA, EGFR (Cat# 2232) Cell Signaling, USA, P-EGFR (1068) (Cat# 2232) Cell Signaling, USA, P-EGFR(1178) (Cat# 2234) Cell Signaling, USA, Erk1/2 (Cat# 9102S) Cell Signaling, USA, P-Erk1/2 (Cat# 9101) Cell Signaling, USA, Anti-ceramide antibody (Cat# C8104) Sigma, USA, GM3 Synthase (ST8SIA-V) (Cat# sc-365329) Santa Cruz, USA, B3GALT4 (Cat# ab169759) Abcam, UK, GM2/GD2 Synthase (B4GALNT1) (Cat# sc-376505) Santa Cruz, USA, Secondary IgG anti-mouse Alexa fluor 594 (Cat# 88903) Cell Signaling, USA, FITC anti-mouse IgM (Cat# 406506) BioLegend, USA, Goat anti-mouse IgG (L+H) (Cat#ab6789) Abcam, UK, Goat anti-rabbit IgG-HRP (Cat# sc-2004) Santa Cruz, USA, Mouse IgG2a Isotype Control (Cat# 02-6200) ThermoFisher, USA.

## METHODS

### Cell culture

Human breast cancer cell lines MCF-7 and BT-474 obtained from American Type Culture Collection (ATCC, Manassas, VA USA) was cultured in DMEM media with 10% Fetal bovine serum, 100 units/mL penicillin and 100µg/mL streptomycin. Cells were grown at 37 ^ᵒ^C with 5% CO_2_ in humidified incubator.

### Generation of MCF-7_RICTOR^SH^ and BT-474_RICTOR^SH^ cell lines

MCF-7 cells were seeded in 48 well plate having DMEM media supplemented with 10% FBS. After 24 h when cells reached 60-70% confluency, media was removed and solution mix containing polybrene, serum free media and lentiviral particles with a combination of shRNA plasmids were added with Multiplicity of Infection (MOI) of 2. The plate was sealed with parafilm, and kept at 37 ^ᵒ^C for 30 min, followed by centrifugation at 1500 rpm at 30 °C for 30 min. The plate was kept back in incubator for 4-6 h with careful monitoring of media to avoid evaporation. After 4-6 h of monitoring, DMEM media supplemented with 20% FBS was added to the cells. The cells were kept with complete media for 24-48 h after which selection with puromycin was performed. Puromycin resistant cells were taken, and expanded for further experiments. MCF-7_RICTOR^SH^ and MCF-7_SCRAM^SH^ cell lines using RICTOR shRNA or scrambled shRNA were generated. Similarly BT-474_RICTOR^SH^ and BT-474_SCRAM^SH^ cell lines were generated.

### Cloning

UGCG and ZFX genes were cloned in pBBL-FLAG vector (BioBharti LifeScience Pvt Ltd) using Xho1 (NEB, R0104S) and Hind III (NEB, R0146S) restriction enzymes. For amplification, the recombinant plasmids were transformed in *Escherichia coli* (DH5α) competent cells, and the positive colonies were selected using kanamycin. The UGCG and ZFX plasmids were extracted using the plasmid extraction kit.

### Transfection Protocol for plasmid DNA/siRNA

MCF-7 and BT-474 cells were transfected with plasmid DNA (UGCG or ZFX or empty vector) or siRNA (targeting UGCG or ZFX or scrambled) using Lipofectamine 2000. About 3.5 x 10^5^ MCF-7/BT-474 cells per well were seeded in six-well plate, in Dulbecco’s Modified Eagle’s Medium-high glucose with 10% FBS and 10% penicillin and streptomycin, and incubated at 37 °C in CO_2_ incubator for 24h. At 80-85% confluency, plasmid and Lipofectamine complex in a 1:3 ratio were incubated for 25 min in Mammary Essential Medium Eagle, and cells were transfected with these complexes. After 6 h of transfection, antibiotic-free media containing 10% FBS was added to the cells, and cells were incubated for 36 h. For generation of stable cell lines, transfected cells were selected with G418 antibiotic resistant marker.

### Collection of patient tumor tissue

Patient tumor and adjoining normal breast tissue were collected from operable breast cancer Luminal patients (Stages I, II, IIIa) undergoing treatment at Dr. B. R. Ambedkar Institute-Rotary Cancer Hospital (BRA-IRCH), All India Institute of Medical Sciences (AIIMS), New Delhi, and from biorepository of Rajiv Gandhi Cancer Institute and Research Center (RGCIRC), Delhi after due ethical approvals. Informed consent was taken from all the patients before acquiring samples. Inclusion criteria included women patients of all age (18–85 years) and socioeconomic status who gave consent for tissue collection, patients with operable breast cancers (Stages I, II, IIIa) who will undergo adjuvant therapy, and patients with ER^+^PR^+^HER2^+^ status. Exclusion criteria included patients who are undergoing neo-adjuvant chemotherapy, and patients who are not fit to undergo surgery. Details of patients and tumor signature are mentioned in **Supplementary Data Set 1.**

### Cellular assays

For proliferation assay, stable cell lines or siRNA (Scrambled siRNA/UGCG siRNA/ZFX siRNA) transfected cells (5000 cells/well) were seeded in a 96 well plate for 24, 48, and 72 h in complete DMEM media, and incubated at 37 °C. Cell proliferation was quantified using MTT assay at 570nm following previously described method (55). For scratch wound migration assay, we followed the previously standardized protocol (55).

To test the effect of different gangliosides on cell proliferation, GD3 and GM1 ganglioside were resuspended in serum-free medium and briefly sonicated for proper nano micelles formation. After 24h, cells were treated with 50 µM of GD3 or GM1 ganglioside and further incubated for 48h for MTT assay.

For transwell migration assay, stable cell lines or siRNA (Scrambled siRNA/UGCG siRNA/ZFX siRNA) transfected cells (60,000 cells) were seeded in DMEM media without FBS in transwell inserts (8 µm pore size), and placed on to 24-well cell culture plates containing DMEM with 10% FBS. After incubating for 24h at 37 °C, non-migrating cells on the upper surface of the membrane were removed with cotton swabs and migrated cells at the base of the inserts were fixed in 4% PFA (∼5 min), permeabilized with methanol (20 min), and stained with 2% crystal violet dye (15 min) followed by PBS washing to remove the extra stain on cell surface. Cells were counted manually and imaged using a microscope.

For anchorage-dependent assay, stable cell lines or siRNA (Scrambled siRNA/UGCG siRNA/ZFX siRNA) transfected cells were seeded in a six-well plate at 2500 cells per well. After 10 days, colonies were washed with 1XPBS, fixed in chilled methanol for 20 min, and stained with 2% crystal violet for 30 min. The wells were imaged using a digital camera, and representative individual colonies or colony forming units comprising >30 cells were counted manually under the stereomicroscope (Leica S8 APO).

### Immunofluorescence in cells

Cells for immunofluorescence were grown on coverslip in complete DMEM medium. After 24h, cells were fixed in 4% paraformaldehyde (PFA) for 20 min, washed with PBS, and were blocked with PBS containing 5% goat serum for 1h at room temperature (RT). The cells were incubated for 2h at 4 °C with appropriate concentrations of respective primary antibodies (RICTOR, Ceramide, UGCG, ZFX), washed thrice with PBS, incubated with the appropriate secondary antibody for 2h at RT, and washed with PBS. The coverslips were then incubated with Hoechst in PBS for 10 min, washed three times thoroughly by gently dipping in distilled water for removal of residual salts of the wash buffer. Coverslips were mounted with ProLong Gold. Images were captured using the Leica TCS SP5 or SP8 confocal microscope using identical settings for a particular staining across treatments for all replicates.

### Pellet collection for RNA and protein isolation

For RNA isolation, cells were grown in 100 mm cell culture plates. For pellet collection, media was aspirated from the plates and washed 2 times with DPBS. 1 mL of Trizol was added to the plate, and incubated for 5 min. After incubation, cells were scraped and transferred into 1.5 mL centrifuge tubes, and used immediately or stored at −80°C. For protein isolation cells were similarly washed, scraped out in DPBS, centrifuged at 5000rpm for 5 min, and used immediately or stored at −80°C.

### Quantitative Real-Time PCR

Total RNA was extracted using already standardized protocol (1). The concentration of the RNA was determined using Nanodrop 2000 (ThermoScientific). The integrity of the RNA was checked on 1% agarose gel. cDNA synthesis and Real-time PCR were done as described previously (55). Relative quantitation of gene expression was done using *β−*actin as the endogenous reference gene for normalization. All primer sequences used for RT PCR are listed in **Supplementary Table 2**.

### Western Blotting

Protein expression analysis was done by western blotting as per previously described method (55). Protein separation was done on 10-12% SDS-PAGE with 25-60 μg of protein. After separation, protein were transferred to the PVDF/Nitrocellulose membrane. Immunostaining was done by overnight incubation of blot with corresponding primary antibody at 4 °C in 5% BSA/Skimmed milk in TBST. After washing, the blots were incubated with secondary antibody for 1 h, and X-ray sheets were developed using Immobilon Western Chemiluminescent HRP.

### Bioinformatic Analysis

Differential gene expression analysis was performed on mouse rictor (-) microarray datasets (GSE46515, GSE67077, GSE84505, GSE85555) obtained from NCBI GEO dataset using the limma package in R (56–60). To study the effect of *RICTOR* knockdown only on the transcription factors (TFs), the downregulated genes were compared with a list of mouse TFs from the Animal TFDB 3.0 database and categorized accordingly (61). Finally, to study whether RICTOR knockdown had any effect on the TFs that bind to the UGCG promoter, the list of transcription factors was then compared with a list of experimentally validated TFs binding to UGCG promoter region (−3kb upstream of ATG) obtained from the TRANSFAC software (62). From this analysis, three potential TFs that may bind to the UGCG promoter were identified.

### ChIP qRT-PCR Primer Designing

For designing the ZFX and H3K4ME3 ChIP primers, we used ChIPBase v2.0 data base (http://rna.sysu.edu.cn/chipbase/). For ZFX primers, we have selected the transcription factor in factor type option, and looked for binding of ZFX transcription factor on UGCG promoter in DAOY medulloblastoma cell line. From the genome of DAOY medulloblastoma cell line, we have selected the ZFX binding region (coordinates; 111896151-111896592) on UGCG promoter provided by ChIPBase v2.0 for designing the ZFX ChIP primer. For H3K4Me3 ChIP primers, we selected the histone modification in factor type option, and looked for binding of H3K4ME3 on UGCG promoter, and selected the region of UGCG DNA (coordinates; 111896051-111898139) for H3K4ME3 ChIP primer. Primer sequences for ChIP PCR are given in **Supplementary Table S2**.

### Chromatin Immunoprecipitation (ChIP)

Cells were grown in 100 mm dish, and after 80-90% confluency (approx. 10-15 million cells), cross linking of the proteins in cells was done using 1% formaldehyde. Formaldehyde was directly added in the dish followed by incubation for 10 min at room temperature, and reaction was quenched by adding 1/8 volume of 1M glycine, and incubated for 5 min. After washing with PBS, cells were scraped in PBS, and resuspended in 1.5 mL nuclear lysis buffer (1% SDS, 10 mM EDTA, 50 mM Tris-HCl pH-8.0, 1X protease inhibitor cocktail). Cells were then sonicated using Bioruptor (Diagenode, Denville, New Jersey, USA) for 38-44 cycles keeping maximum amplitude with 30 sec pulse and 30 sec hold. Chromatin shearing was checked on 1% agarose gel to confirm the correct size (approximately 300-500bp) of chromatin. Protein estimation were done using Bicinchoninic acid (BCA) protein estimation kit according to manufacturer protocol. Preclearing was done by incubating protein with A/G magnetic beads for 1h at 4 °C while rotating. For immunoprecipitation, 500 µg of the chromatin was incubated with 2.5 µg of antibody (ZFX, H3, H3K4Me3, or IgG) overnight at 4 °C while rotating. Next day, 50 µL of protein A/G magnetic beads were added, and incubated for an additional 2 h. Each sample was then subjected to one wash with 1 mL of low salt buffer (0.1% SDS w/v, 1% Triton X 100, 2 mM EDTA pH 8.0, 150 mM NaCl), high salt buffer (0.1% w/v SDS, 1% Triton X 100, 2 mM EDTA pH 8.0, 500mM NaCl), LiCl buffer (20mM Tris HCl, 500nM NaCl, 2mM EDTA pH 8.0, 0.1% w/v SDS, v/v 1% IGEPAL), and finally 3 washes with TE buffer (10 mM Tris HCl pH 8.0, 1 mM EDTA pH 8.0). All washes were incubated for 5 min at 4 °C while rotating. Next, 500 µL elution buffer (1M NaHCO3, 10%SDS) was added for elution of chromatin from antibody followed by incubation for 5 min. Reverse crosslinking was done by adding 4 μL of 10% SDS (final concentration, 0.2%) and incubating at 65 °C overnight in a mixer with constant agitation at 1200 rpm. RNase treatment was done for all the samples (including the input samples) by adding RNase A (100µg/mL) to a final concentration of 0.2 µg/µL, followed by incubation for 2 h at 37°C. Proteinase K (10 mg/ml) was added to the samples to a final concentration of 200 µg/ml followed by incubation of 2 h. The DNA isolation was done by phenol/chloroform/isoamyl alcohol (25:24:1) method. The DNA pellets were air dried, and finally dissolved in 20 µL of 10 mM Tris buffer (pH 8.0). Final DNA estimation was done using Qubit dsDNA HS assay kit. ChIP-qPCR was carried out with equal amount of ChIP DNA per reaction.

### ChIP RT-PCR

For ZFX ChIP RT-PCR the ChIP DNA was quantified using Qubit HS DNA Kit, and ChIP-qPCR was performed with equal amount of DNA from ChIP, Input and its IgG control DNA. The mean Ct value was used to calculate fold change after normalization with IgG and Input. For H3K4ME3, the normalization was done using input and total H3. For ZFX peak calling, ENCODE (PMID: 22955616; PMCID: PMC3439153, PMID: 29126249; PMCID: PMC5753278) database was used.

The ENCODE identifiers used was ENCSR435OQD (ZFX). For both of the transcription factors, the peaks were visualized using the UCSC genome browser.

### Isolation and quantification of sphingolipids. using LC-MS/MS

Collection of cell pellets, lipid isolation, LC-MS/MS analysis, and absolute quantitation of sphingolipids were done as per published protocol (55).

### Isolation and quantification of gangliosides using LC-MS/MS

Lipids were extracted from ∼15 × 10^6^ cells. Cells were washed in ice-cold PBS, harvested, and centrifuged. The cell pellet were resuspended in methanol (200 µL), and homogenized by sonication. An aliquot (5 µL) was taken for protein estimation by Bicinchoninic acid (BCA) protein estimation kit. 3 mg protein equivalent of each sample were used for lipid isolation. The cell suspension were transferred to Teflon-lined borosilicate tubes containing 3 mL chloroform : methanol (2:1) and deuterated monosialoganglioside, GM3-d3 (d18:1/18:0-d3) internal standard. Samples were sonicated and centrifuged, and supernatant was transferred to fresh tubes. The residue were re-extracted with 3 mL chloroform : methanol (1:1), and a third time with 2.5 mL chloroform : methanol : water (1:2:0.8), and all supernatants were pooled. 200 µL of 0.1M sodium hydroxide (NaOH) was added to the pooled extract, and incubated with shaking at 37 °C for 2 h. The solvents were then evaporated under N_2_ gas, and resuspended in methanol : water (1:1). For enrichment of gangliosides, samples were passed through C18 Sep-Pak cartridge. The Sep pak column was washed with 5 mL methanol, followed by 1 mL chloroform : methanol (1:1), 1 mL water, and finally equilibrated with methanol : water (1:1). Sample was passed through the Sep-Pak column. Elution was done with 2 mL methanol : water (1:1), 1 mL methanol, 1mL methanol : chloroform (1:1, v/v) and finally with 1mL chloroform : methanol (2:1). All the eluents were pooled, and evaporated under N_2_ gas. Dried samples were resuspended in 200 µL of solvent B [methanol : isopropanol (1:1) with 0.2% formic acid and 5 mM ammonium acetate)]. The samples were vortexed, centrifuged, and transferred to the autoinjector vial for LC-MS/MS analysis using high pressure UHPLC liquid chromatograph (Exion LC AC, SCIEX, USA) coupled to a hybrid triple quadrupole/ linear ion trap mass spectrometer (4500 QTRAP, SCIEX, USA). A Kinetex® C18, 2.1 × 50 mm column (Phenomenex®) with a particle size of 1.7 µm was used at oven temperature of 55 °C. Total optimized run time was 20 min where solvent A (2% methanol, 0.2% formic acid and 5 mM ammonium acetate in water) and solvent B [methanol : isopropanol (1:1) with 0.2% formic acid and 5 mM ammonium acetate)] were used as mobile phase A and B, respectively with a gradient flow rate of 0.3 mL/minute from 0 to 7 minutes and then changes to 0.4 mL/minute for total run followed by initial conditions in the next run. The LC retention time was noted for each analyte. The sample dilution were standardized by confirming the peak profile and intensity for each transition at the retention time. The analyte peak area for all experiments were integrated in MultiQuant 3.0.2 for data analysis. The LC gradient buffer flow system of solvent A and solvent B were maintained at a ratio of solvent A : B (60:40) for 2 min, then ratio (30:70) for 5 min, followed by a linear gradient of (5:95) from 7 to 13.5 min then washed with solvent A : B (60:40) for 13.5 to 20 min before subsequent run. Ganglioside estimations were done using multiple reaction monitoring (MRM) with Enhanced Product Ion (EPI) scan. Collision cell exit potential (CXP) and Collision energy (CE) were optimized. Source dependent parameters like nebulizer gases GS1 (40psi), GS2 (40psi), Curtain gas (35psi), temperature (600°C), ion spray voltage (−4300 KV) and CAD (high) were optimized. A standard curve was generated for absolute quantitation of gangliosides. To quantify absolute value of analyte in each sample, the area under the peaks for both analyte and spiked internal standard (GM3-d3) was estimated, and ratio of each analyte with internal standard was determined. These ratio values were used to quantitate all ganglioside species in cell extracts. Three independent biological replicates (n = 3) were used, and for each sample, three technical replicates were run with two blank runs between each sample.

### Flow cytometry for gangliosides

Cells were seeded in 12 well plates and pellets were collected after 72 h incubation. Cells were then washed twice with staining buffer (0.5% BSA in 1 X PBS). For GD3 gangliosides, cells were incubated with anti-GD3 (1:50) for 1 h on ice. After washing, cells were stained for 1 h on ice with Alexa fluor 488 anti-IgG (1:50). Cells were then washed by PBS-BSA buffer and acquired using BD FACSVerse. Control experiments were also performed with secondary antibody alone. For GM1 ganglioside, we used FITC labelled cholera toxin B (1 μg/test) for 30 min on ice. Data was analysed using Flowjo.

### Animal Studies

All tumor growth kinetic studies were performed using MCF-7 and its derivatized cell lines (MCF-7_SCRAM^SH^, MCF-7_RICTOR^SH^, MCF-7_VECT^OE^, MCF-7_UGCG^OE^, MCF-7_ZFX^OE^) in NOD SCID C.B-17 mice. The flank region of the mice were shaved with Veet to remove the hair before the cell injection. The cells, suspended in FBS:Matrigel (1:1, 200 µL), were then injected (1.5*10^6^) subcutaneously in the flank region. Once the tumor volume reached 40-50 mm^3^, the tumor volume and body weight of the mice was measured after every 2 days and ten readings were taken for every experiment unless stated otherwise. Tumor volume was calculated using formula L*B^2^/2 where L is length and B is breadth. On final day of the measurement, the tumor were harvested, washed with 1X PBS, and cut into two halves. One half was stored in 4% PFA for immunofluorescence imaging, and other half was stored in Allprotect tissue reagent at −80°C for molecular studies.

We conducted independent experiments to compare the tumor growth kinetics of MCF-7_ SCRAM^SH^ and MCF-7_RICTOR^SH^, or MCF-7_ VECT^OE^ and MCF-7_UGCG^OE^, or MCF-7_ VECT^OE^ and MCF-7_ZFX^OE^.

For siRNA experiments, mice were randomized into different groups after the tumor reached 50 mm^3^ volume, group 1 mice were left untreated, group 2 mice were treated with scrambled siRNA, group 3 mice were treated with target siRNA. Group 2 and 3 mice were treated with 50 µL of 200 ng of siRNA complexed with TAC6 polymer (siRNA: TAC6 polymer; 1:10) at the tumor site. A total of 6 doses were given at an interval of every two day (63, 64). At the end of the experiment tumor was excised and stored at −80° C for further analysis.

For inhibitor experiments, mice were randomized into different groups after tumor reached 50 mm^3^ volume. Group 1 mice were left untreated. In group 2, mice were implanted with blank hydrogel, and group 3, 4 mice were implanted with hydrogel entrapping KDM5A inhibitor with a total dose of 2.5 mg/kg and 5 mg/kg respectively (27). After every 2-day tumor volume was measured for different groups.

### Immunofluorescence in tissue sections

Patient tumor and adjacent normal tissue frozen at −80 °C in Allprotect tissue reagent were processed and fixed and stained following already described method (55). The tissue sections were stained with RICTOR, Ki67, UGCG, ZFX and anti-ceramide antibodies. Confocal imaging of the samples was performed with Leica TCS SP8 microscope. The sections were visualized at 40X oil immersion using LAS AF software. Z-stacking was performed, and images were acquired for each section. The images were processed using LAS X software.

### Immunohistochemistry (IHC) in human tissue samples

IHC for UGCG and ZFX was carried out on Tissue micro array sections of Breast cancer using standard protocol after due ethical approval. Briefly, 5 µm FFPE tissue sections were fixed on glass slides, followed by deparaffinization in xylene, rehydration in graded alcohol, 3% H2O2 treatment for 30 min, antigen retrieval using citrate buffer (pH 6.00) for 15 min, and blocking with 3% bovine serum albumin for 30 min. After this, sections were incubated with primary antibody against UGCG (dilution 1:250) and ZFX (dilution 1:100) for 1h at room temperature. Incubation with secondary antibody (DAKO REAL™EnVision™) was done for 30 min. 3-3’ Diaminobenzidine (DAB, DAKO REALTMEnVisionTM) for 10 min was used as chromogenic substrate, and haematoxylin was used as a counterstain. Sections were dehydrated, dried, mounted with DPX, and visualized under microscope. All sections were examined by pathologist to score tumor cells. Sub-localization of the staining in the cytoplasm and nucleus was recorded separately. Clinical details of patients is provided in Supplementary Data set S2.

### Analysis of the TCGA and METABRIC datasets

We accessed the TCGA dataset (https://www.cancer.gov/tcga). This dataset had RNS sequencing data of 1082 breast cancer patients that were available. Similarly, METABRIC data set was accessed with 1905 tumors with gene expression levels available from microarray (65). In both datasets, expression levels of UGCG and ZFX were compared between PAM 50 subtypes and by estrogen receptor status. Total of 601 tumors were identified as Luminal A+B after elimination of other tumors within TCGA and 1170 within METABRIC datasets. Mean expression level of UGCG and ZFX was used as a cut-off within the luminal tumors to divide the tumors as high and low for both genes respectively. Expression levels of ZFX were compared between the high and low UGCG expressing tumors within the luminal groups.

### RNA Isolation & Quantitative Real-Time PCR from patient tumor tissue

RNA isolation and quantitative RT PCR from Luminal A Patient tissue samples (∼20 mg) stored in Allprotect Tissue Reagent were performed using already reported method (55). cDNA synthesis and Real-Time PCR were done as described above.

### Statistical Analysis

Statistical analyses were carried out using Graph-Pad Prism 7 (GraphPad Software). All data are represented as mean ± SD or mean ± SEM. A minimum of three or more biological replicates were used per condition in each experiment as mentioned in each Figure legend. Pairwise comparisons were determined using Student’s t-test. Multiple comparisons among groups were determined using one-way ANOVA followed by a post-hoc test. Growth kinetic analysis were performed using two-way ANOVA. Differences between groups were considered significant at *p*-values below 0.05 (\**p* < 0.05, \*\**p* < 0.001, *** *p* < 0.0001, and \*\*\*\**p* < 0.0001).

## ACKNOWLEDGEMENTS

We are grateful to Dr. Sagar Sengupta, Dr. Vinay Nandicoori, Dr. Kaustav Bandopadhyay, and Dr. Tapas Mukherjee for many helpful discussions. pERK1/2 antibody was kind gift from Dr. Vinay Nandicoori. We thank DST-FIST sponsored Amity Lipidomics Research Facility at Amity University Haryana. We thank biorepository of Rajiv Gandhi Cancer Institute and Research Centre (RGCIRC), Delhi. Figure 7 was drawn using BioRender.com with publication licence numbers OZ23FCUP9X and PA23FCUPHN.

## FUNDING

The support from Amity University Haryana, RCB, AIIMs, and Department of Biotechnology (DBT) Govt. of India is greatly acknowledged. Research in U.D. group is supported by BT/PR19624/BIC/101/488/2016 (DBT), BT/PR40413/BRB/10/1922/2020 (DBT), CRG/2021/002966 (SERB), and 5/13/81/2020/NCDIII (ICMR). Research in AB group is supported by BT/PR40413/BRB/10/1922/2020 (DBT). AM is supported by SERB-STAR award (STR/2019/000064) and DBT-National Bioscience Award for Career Development (BT/HRD/NBA/38/04/2016). Amity Lipidomics Research Facility at Amity University Haryana is supported by DST-FIST grant, SR/FST/LSI-664/2016. M.N.A., T.P. and A.K. thank ICMR, and S.J. and K. Rana thank CSIR for research fellowships. We are grateful to Nadathur Estates for their support of all the breast cancer research activities at SJRI. We acknowledge the support of the DBT e-Library Consortium (DeLCON) for providing access to e-resources. Animal work in the small animal facility of Regional Centre for Biotechnology is supported by BT/PR5480/INF/22/158/2012 (DBT).

## ETHICAL STATEMENT

All animal experiments were performed after due approval of the Institutional Animal Ethical Committee of Regional Centre for Biotechnology (RCB/IAEC/2020/079) as per the guidelines of Committee for the Purpose of Control and Supervision of Experiments on Animals (CPCSEA), India. All studies with human tissue samples were conducted after due ethical clearance from AIIMS (IEC-332/01.07.2016), RGCIRC (RGCIRC/IRB/276/2019, Res/BR/TRB-20/2020/ 70), Amity University Haryana (IEC-AIISH/AUH/ 2016-1), and SJRTI (S475/79-80).

## AUTHOR CONTRIBUTIONS

K. Rajput performed all cell culture experiments and immunofluorescence studies. M.N.A. performed and analysed all gene and protein expression studies and ChIP analysis. S.K.J. and K.Rajput performed the animal studies. N.M., P.S., K.C., and A.K. performed sphingolipid and ganglioside estimation experiments. S.D performed the bioinformatic analysis. K. Rana performed flow cytometry experiments. A.K. created shRNA silenced cell lines, and performed activity assays. T.P., G.M, and J.P. compiled patient clinical information. S. Deo provided Luminal patient tissue samples, and patient clinical information. J.P. performed the immunohistochemistry analysis and metadata analysis. A.M. supervised bioinformatic analysis and validation. K.R., M.N.A., A.K. compiled the data. All authors read the manuscript and gave their input. U.D. conceived the idea. U.D. and A.B. wrote the manuscript, and supervised the whole project.

## SUPPORTING INFORMATION

Complete list of supporting figures, tables, and data sets are in supporting information.

## Notes

### Competing Interest Statement

The authors have declared no competing interest.

### Summary of Updates

We validated our studies in another breast cancer cell line (BT-474 cell line) that represents luminal B breast cancer subtype, and revised the manuscript extensively. Now, our results clearly demonstrate the molecular mechanism and signalling circuitry of mTOR/RICTOR regulating ganglioside metabolism in two luminal breast cancer cell lines (MCF-7 and BT-474) supported by murine data. To further validate if ZFX-mediated UGCG regulation is a common phenomenon among other cancer cell types, we showed that ZFX silencing downregulates UGCG expression in other cancer cell types like HCT-116 (colon cancer), HepG2 (liver cancer), and MDA-MB-453 (HER2+ breast cancer representative). This has added the dimension of robustness to our already validated metabolic-signalling-gene regulatory circuit in breast cancer subtypes. In addition, we also performed the validation/rescue studies using gangliosides. In the revised manuscript, we show that addition of GD3 gangliosides on MCF-7_RICTORSH and BT-474_RICTORSH cells enhances the cell proliferation, and addition of GM1 gangliosides on MCF-7_UGCGOE, MCF-7_ZFXOE, BT-474_UGCGOE, and BT-474_ZFXOE cells decreases cell proliferation. This provides more confidence to the existing data that has established the circuit connecting mTORC2/RICTOR to ganglioside metabolism controlling tumor progression. We have included these results in the revised manuscript.

