## Supplementary Information for "RICTOR Drives ZFX-mediated Ganglioside Biosynthesis to Promote Breast Cancer Progression"

##### Table of Contents


| S. No. | Item | Page No |
|---|---|---|
| 1. | Figures S1-S6 | S2-S12 |
| 2. | Table S1 | S13 |
| 3. | Table S2 | S14 |
| 4. | Table S3 | S15 |
| 5. | Legends for data sets 1-3 | S16 |

#### Supplementary Figure 1

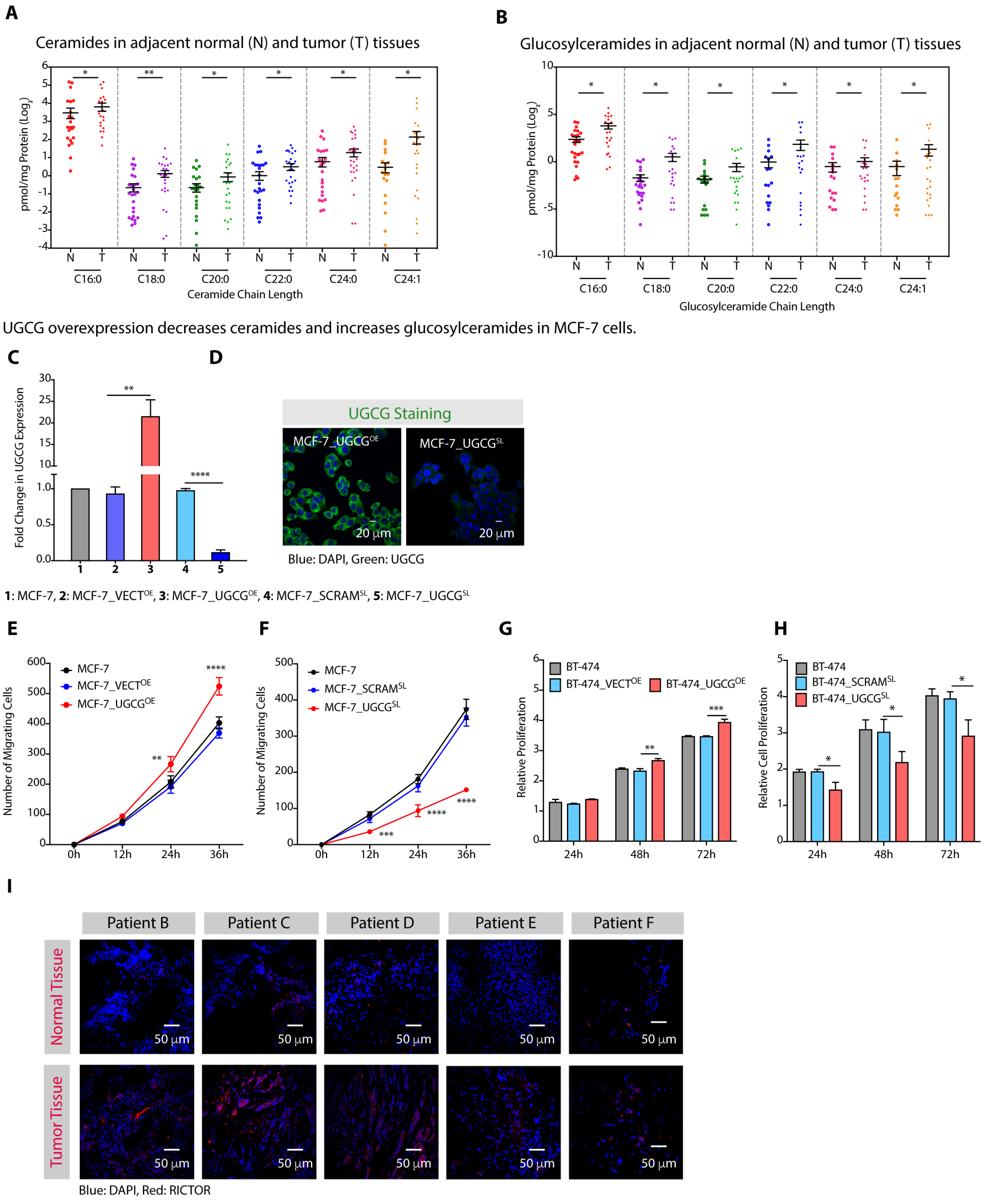

**Supplementary Figure S1. (A, B)** Absolute quantitation (pmol/mg protein) of different species (C16:0, C18:0, C20:0, C22:0, C24:0, and C24:1) of ceramides **(A)** and glucosylceramides **(B)** show higher levels in luminal tumor tissues (labelled as T) in comparision to adjacent normal tissues (labelled as N). **(C, D)** qRT-PCR (mean ± SD, n = 6) **(C)** and immunofluorescence **(D)** validations showing an increase in UGCG expression in MCF-7_UGCG$^{OE}$ cells and effective silencing of UGCG in MCF-7_UGCG$^{SL}$ cells. **(E, F)** Scratch wound migration assay (mean ± SD, n = 5) demonstrate an increase in number of MCF-7_UGCG$^{OE}$ migrating cells **(E)** whereas MCF-7_UGCG$^{SL}$ cells show significantly reduced number of migrating cells **(F)**. **(G, H)** Cell proliferation (mean ± SD, n = 3) assay confirm an increase in proliferation of BT-474_UGCG$^{OE}$ cells (**G**) whereas BT-474_UGCG$^{SL}$ cells show significantly reduced cell proliferation (**H**). **(I)** Immunofluorescence images show elevated RICTOR expression in tumor tissues as compared to adjacent normal tissue sections. Data in Figure S1A-S1C were analysed using an unpaired student's *t* -test, and in Figure S1E-S1H were analysed using Two-way ANOVA. *p*-value: **p* < 0. 05, ***p* < 0.01, ****p* < 0.001, *****p* < 0.0001.

### Supplementary Figure 2

RICTOR silencing inhibits cell proliferation, migration and invasion in MCF-7 cells.

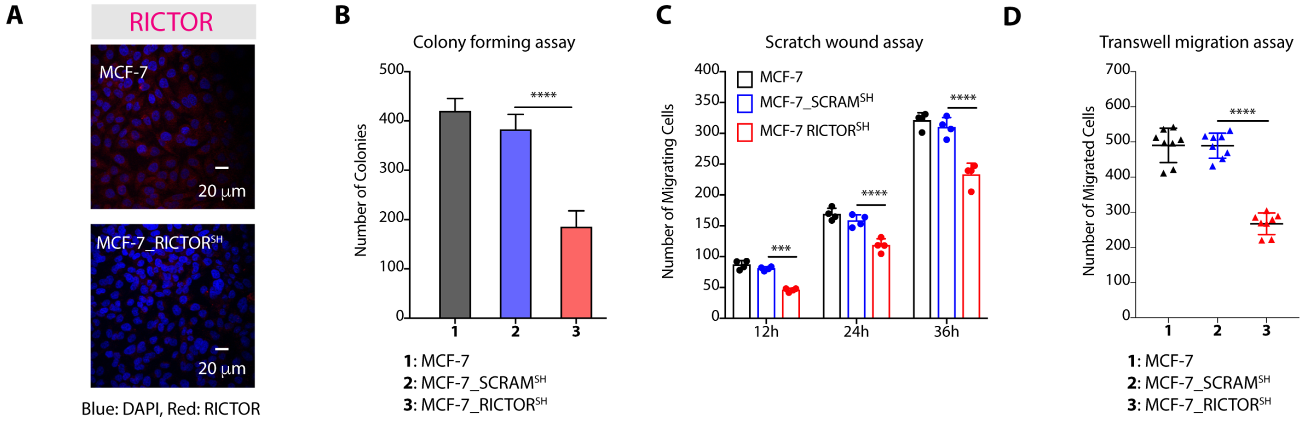


RICTOR silencing inhibits cell proliferation in BT-474 cells via UGCG-mediated regulation of glucosylceramides.

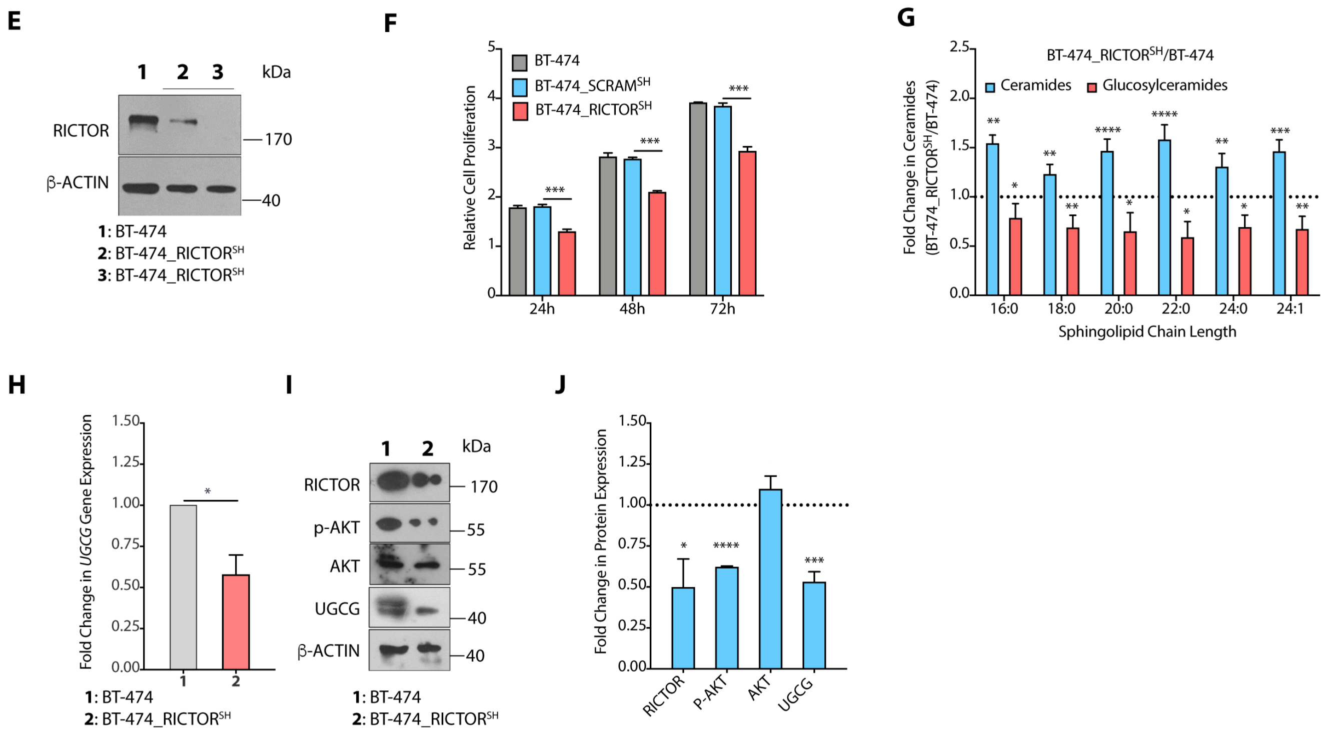

**Supplementary Figure S2.** **(A)** Immunofluorescence staining confirm knockdown of RICTOR. **(B-D)** Colony forming assay (mean ± SD, n = 4) **(B)**, scratch wound assay (mean ± SD, n = 4) **(C)**, and transwell migration assay (mean ± SD, n = 4) **(D)** show decrease in number of colonies and migrating cells on RICTOR knockdown. **(E)** Immunoblots with representative β-actin as control confirm knockdown of RICTOR expression in BT-474_RICTOR$^{SH}$ cells. **(F)** Cell proliferation kinetic studies show decrease in proliferation of BT-474_RICTOR$^{SH}$ cells (mean ± SD, n = 4) as compared to BT-474_SCRAM$^{SH}$ cells. **(G)** Fold change (mean, n = 5) in ceramide and glucosylceramide levels (C16:0, C18:0, C20:0, C22:0, C24:0, and C24:1) in BT-474_RICTOR$^{SH}$ cells as compared to BT-474 cells. **(H, I, J)** qRT-PCR (mean ± SD, n = 3) **(H)**, immunoblots with representative β-actin as control **(I)**, and their quantification (mean ± SD, n = 3) **(J)** demonstrate downregulation of UGCG expression in BT-474_RICTOR$^{SH}$ cells as compared to BT-474 cells. Details of all Immunoblots with originals are provided in supplementary information. Data in Figure S2B, S2D, S2G, S2H and 2J were analysed using an unpaired student's *t*-test, and in Figure S2C and S2F was analysed using Two-way ANOVA. *p*-value: $*p < 0.05$, $**p < 0.01$, $***p < 0.005$, $****p < 0.0005$.

#### Supplementary Figure 3

ZFX alters proliferation, migration, and invasion in MCF-7 and BT-474 cells.

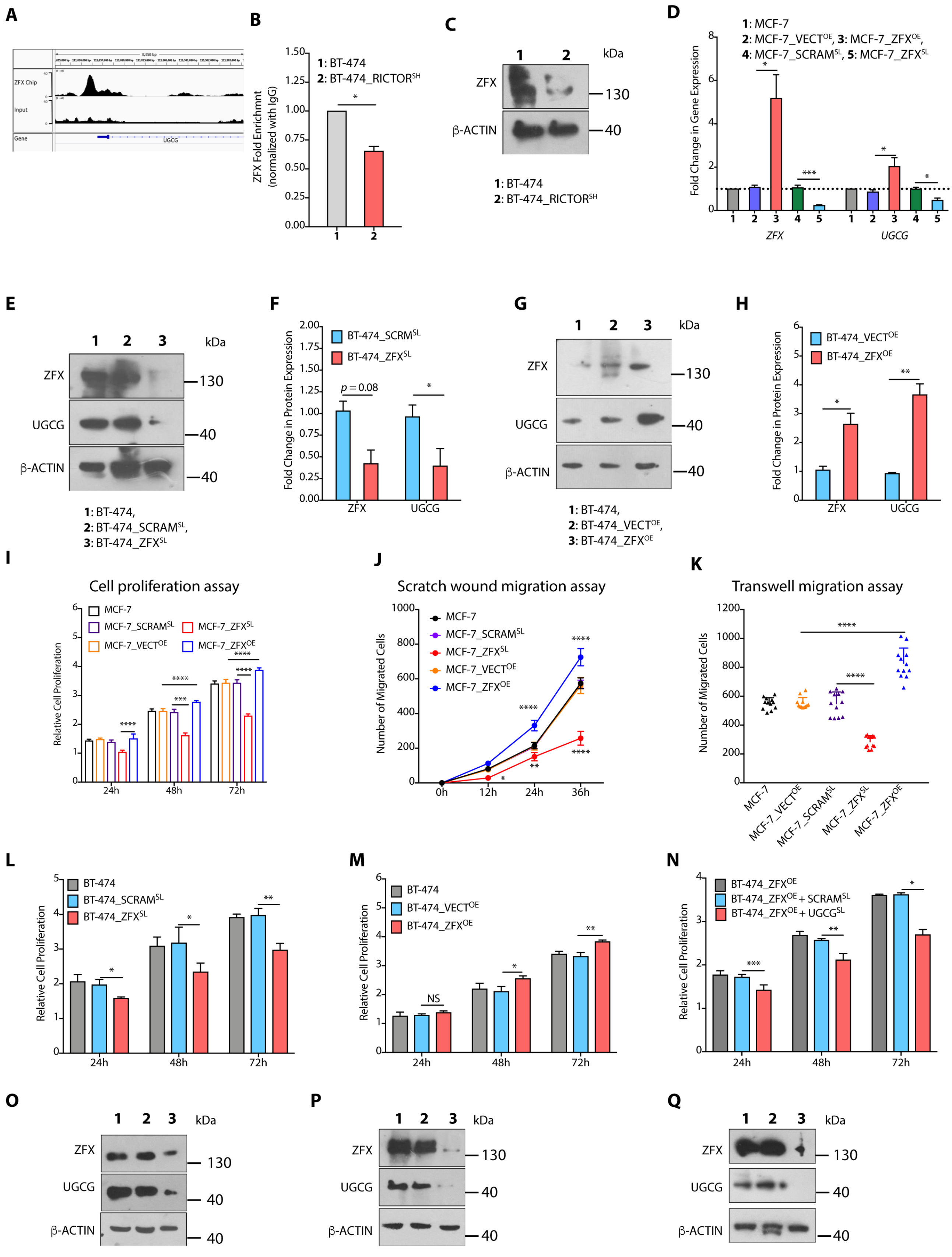


1: HCT-116, 2: HCT-116_SCRAMSL, 3: HCT-116_ZFXSL    1: HepG2, 2: HepG2_SCRAMSL, 3: HepG2_ZFXSL    1: MDA-MB-453, 2: MDA-MB-453_SCRAMSL, 3: MDA-MB-453_ZFXSL

**Supplementary Figure S3. (A)** ZFX binding to UGCG promoter by ZFX ChIP seq. **(B)** ChIP-qPCR (mean ± SD, n = 3) results show reduced binding of ZFX to UGCG promoter in MCF-7_RICTOR$^{SH}$ cells. **(C)** Immunoblot with representative β-actin as control shows downregulation of ZFX expression in BT-474_RICTOR$^{SH}$ cells. **(D)** qRT-PCR (mean ± SD, n = 3) results validate increase in expression of ZFX in MCF-7_ZFX$^{OE}$ cells and silencing of ZFX in MCF-7_ZFX$^{SL}$ cells that results in significant increase in *UGCG* expression in MCF-7_ZFX$^{OE}$ cells, and downregulation of *UGCG* in MCF-7_ZFX$^{SL}$ cells. **(E-H)** Immunoblots with representative β-actin as control **(E, G)** and their quantification (mean ± SD, n = 3) **(F, H)** confirm that silencing of ZFX decrease UGCG expression (**E, F**) and overexpression of ZFX enhances UGCG expression in BT-474 cells (**G, H**). **(I-K)** Cell proliferation (mean ± SD, n = 4) **(I)**, scratch wound assay (mean ± SD, n = 4) **(J)**, and transwell migration assay (mean ± SD, n = 4) **(K)** demonstrate increase in proliferation and migration of MCF-7_ZFX$^{OE}$ cells whereas MCF-7_ZFX$^{SL}$ cells show reduced cell proliferation and migration. **(L, M)** Cell proliferation assay (mean ± SD, n = 4) confirm decrease in cell proliferation of BT-474_ZFX$^{SL}$ cells **(L)** whereas BT-474_ZFX$^{OE}$ cells show enhanced cell proliferation **(M)**. **(N)** siRNA-mediated silencing of UGCG results in significant decrease in proliferation of BT-474_ZFX$^{OE}$ cells. **(O-Q)** Immunoblots with representative β-actin as control confirm that silencing of ZFX decrease UGCG expression in HCT-116 **(O)**, HepG2 **(P)** and MDA-MB-453 **(Q)** cells. Details of all Immunoblots with originals are provided in supplementary information. Data in Figure S3B, S3D, S3F, S3H and S3K were analysed using an unpaired student's *t* -test, and in Figure S3I, S3J, and S3L-S3N were analysed using Two-way ANOVA. *p*-value: *$p < 0.05$, **$p <$ 0. 005, ***$p < 0.001$, ****$p < 0.0005$.

#### Supplementary Figure 4

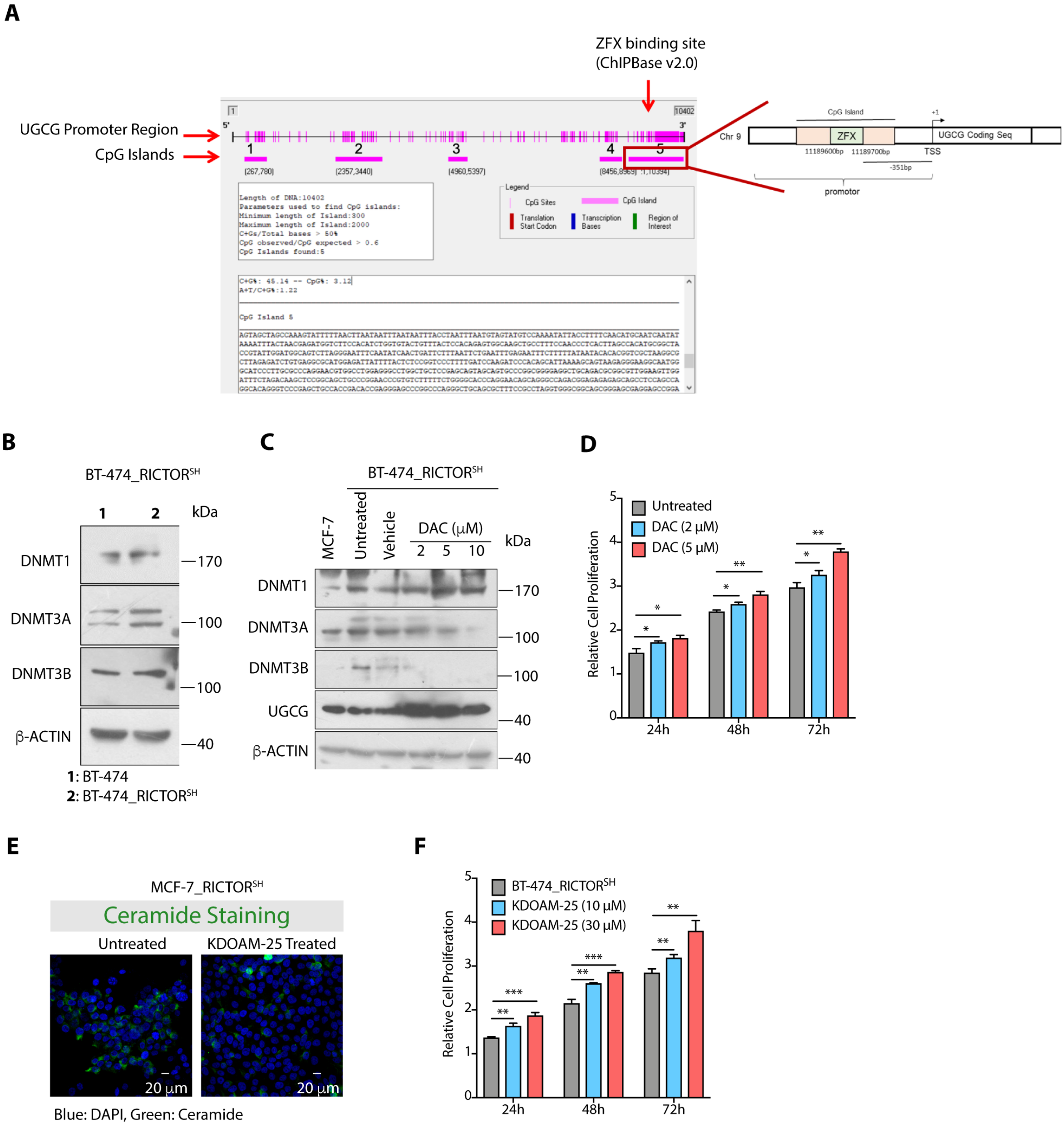

**Supplementary Figure S4. (A)** UGCG promoter site showing multiple CpG islands, and ZFX binding site. **(B)** Immunoblots with representative β-actin as control show an increase in expression of DNMTs in BT-474_RICTOR$^{SH}$ cells as compared to BT-474 cells. **(C)** Immunoblots with representative β-actin as control reveal a decrease in expression of DNMT3A and DNMT3B with a concurrent increase in UGCG expression in BT-474_RICTOR$^{SH}$ cells upon DAC treatment. **(D)** Cell proliferation (mean ± SD, n = 3) assay confirm increase in proliferation rate of BT-474_RICTOR$^{SH}$ cells on DAC treatment. **(E)** Immunofluorescence images confirm enhanced expression of ceramides in MCF-7_RICTOR$^{SH}$ cells on KDM5A inhibition (30 μM). **(F)** Cell proliferation (mean ± SD, n = 3) assay confirm increased proliferation of BT-474_RICTOR$^{SH}$ cells on KDM5A inhibition. Details of all Immunoblots with originals are provided in supplementary information. Data in Figure S4D and S4F were analysed using Two-way ANOVA. *p*-value: **p* < 0.05, ***p* < 0. 005, *****p* < 0.0005.

#### Supplementary Figure 5

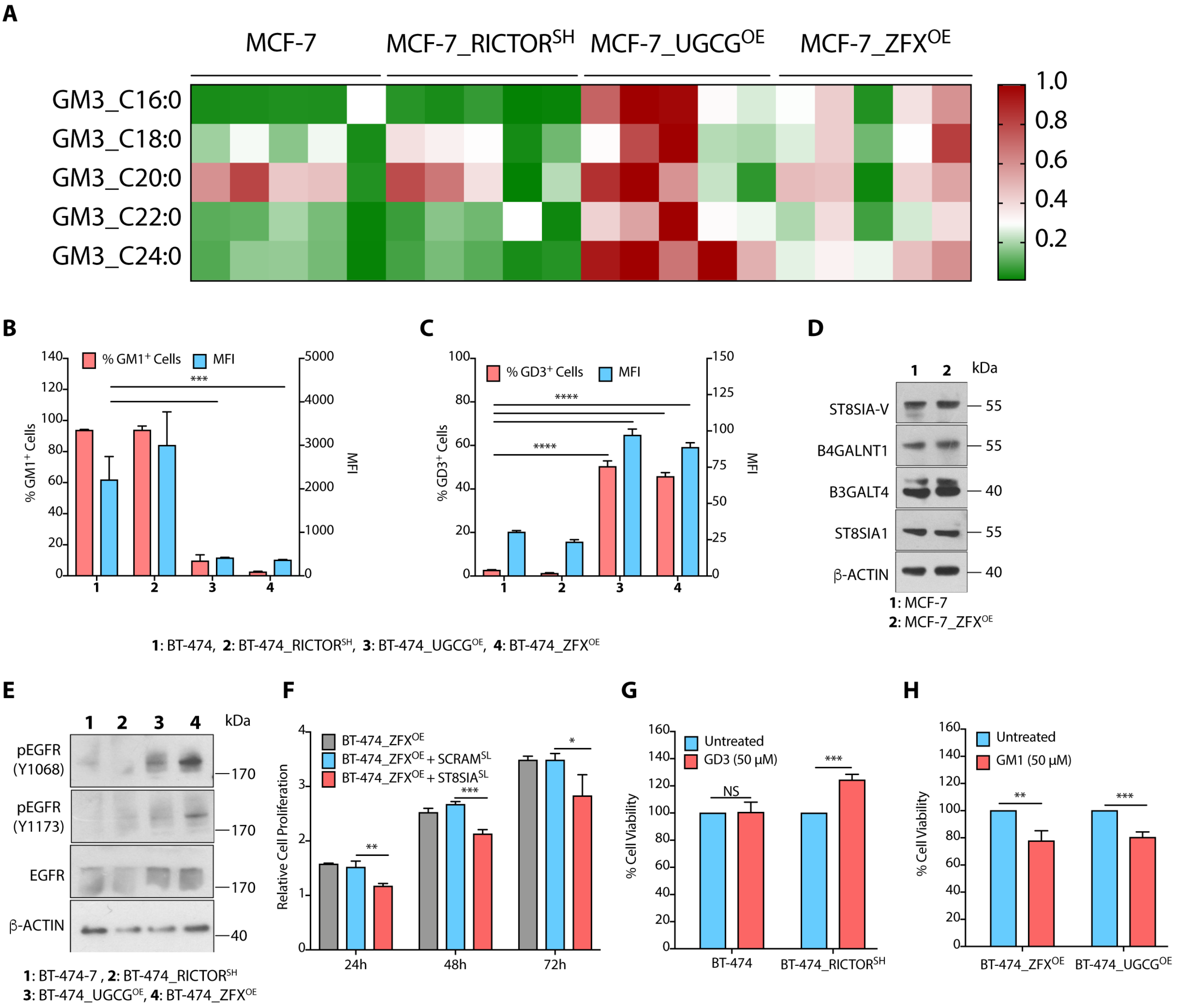

**Supplementary Figure S5.** **(A)** Heat map for normalized quantity (mean, n = 5) (pmol/mg protein) of different species of GM3 (C16:0, C18:0, C20:0, C22:0, C24:0) gangliosides reveal higher levels in MCF-7_UGCG$^{OE}$ and MCF-7_ZFX$^{OE}$ cells as compared to MCF-7 and in MCF-7_RICTOR$^{SH}$ cells. **(B, C)** Quantification of GM1$^{+}$ **(B)** and GD3$^{+}$ **(C)** cells and MFI (mean fluorescence intensity) by flow cytometry confirm decrease in GM1 expression and enhanced GD3 expression in BT-474_UGCG$^{OE}$ and BT-474_ZFX$^{OE}$ cells. **(D)** Immunoblots with representative β-actin as control showing no significant changes in expression of B3GALT4, ST8SIA1, B4GALNT1, and ST8SIA-V in MCF-7_ZFX$^{OE}$ cells. **(E)** Immunoblots with representative β-actin as control reveal an increase in levels of pEGFR$^{Y1173}$ and pEGFR$^{Y1068}$ in BT-474_UGCG$^{OE}$ and BT-474-7_ZFX$^{OE}$ cells as compared to BT-474 cells. **(F)** Cell proliferation assay demonstrates decrease in cell proliferation (mean ± SD, n = 3) of BT-474_ZFX$^{OE}$ cells on siRNA-mediated inhibition of GD3 synthase (ST8SIA1). **(G, H)** Cell proliferation assay show increase in proliferation of BT-474_RICTOR$^{SH}$ cells upon feeding with GD3 gangliosides **(G)**, and decrease in cell proliferation of BT-474_UGCG$^{OE}$ and BT-474_ZFX$^{OE}$ upon feeding with GM1 gangliosides **(H)**. Details of all Immunoblots with originals are provided in supplementary information. Data in Figure S5B, S5C, S5G and S5H were analysed using an unpaired student's *t* -test, and in Figure S3F was analysed using Two-way ANOVA. *p*-value: $^{*}p$ < 0. 05, $^{***}p$ < 0.001, $^{****}p$ < 0.0005.

**Supplementary Figure 6**

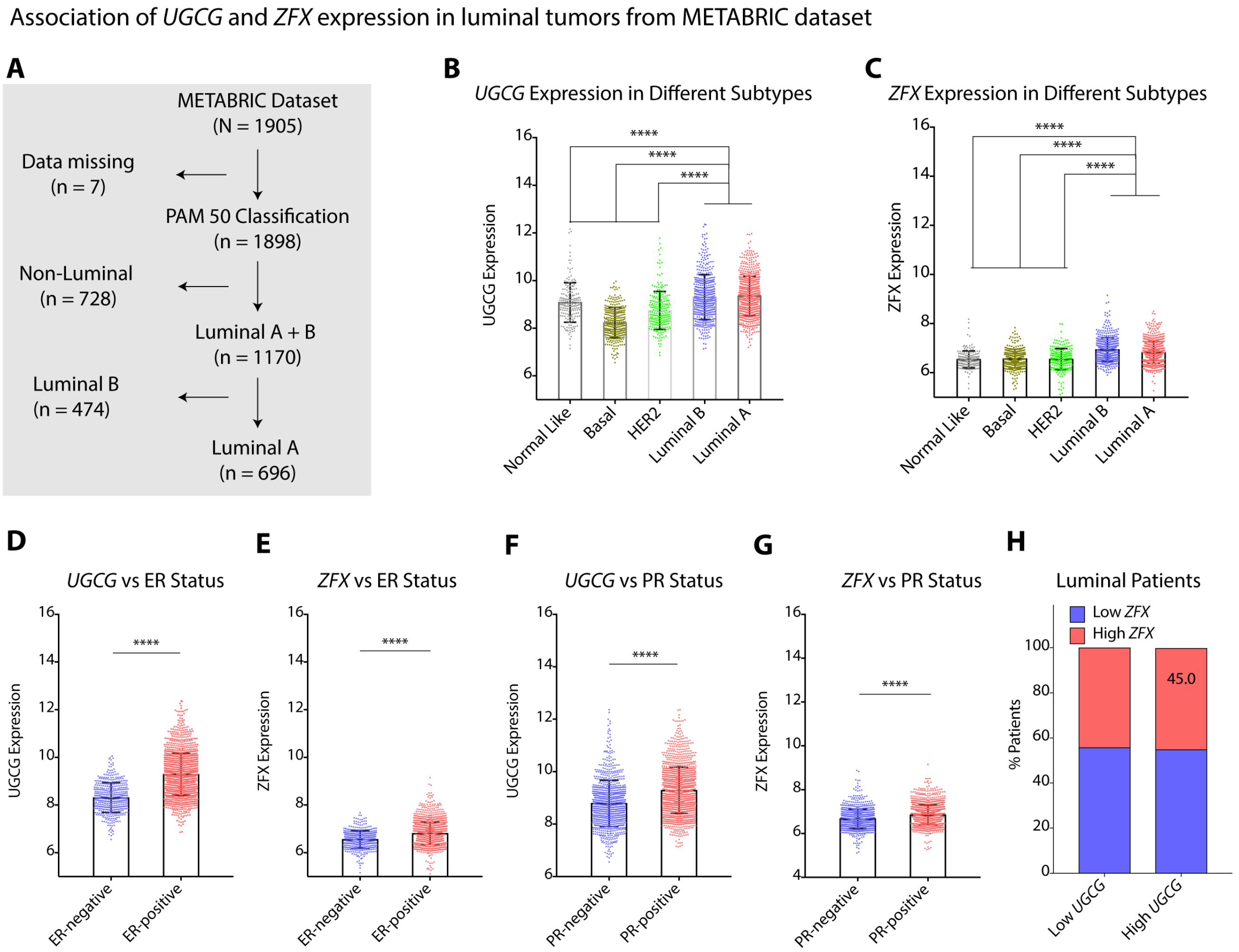


**Supplementary Figure S6. Strong Association of UGCG and ZFX Expression in Luminal Tumors from METABRIC Dataset (A)** Schematic showing the PAM50 classification of METABRIC tumor dataset used for analysis. **(B, C)** Gene expression of *UGCG* **(B)** and *ZFX* **(C)** in different breast cancer subtypes (PAM50) of METABRIC dataset confirm high expression of *UGCG* and *ZFX* in luminal subtypes as compared to other subtypes. **(D-G)** Change in expression of *UGCG* **(D, F)** and *ZFX* **(E, G)** with respect to ER **(D, E)** and PR **(F, G)** status in breast tumors of METABRIC dataset confirm high *UGCG* **(D, F)** and high *ZFX* **(E, G)** expression in $ER^+$ and $PR^+$ tumors. **(H)** Percentage of tumors having high expression of *UGCG* and *ZFX* among Luminal subtype tumors in METABRIC data set. Data in Figure S6B, 6C, S6D-S6G were analysed by an unpaired student's *t*-test. *p*-value: ****$p < 0.0001$.

**Supplementary Table S1.** List of primers (human) used for validation of endogenous gene expression by Real-Time PCR.

| Gene Name | Forward primer (5' to 3') | Reverse primer (5' to 3') |
|---|---|---|
| *CerS1* | GTCATGTGGCGCCTGTTTC | CCAGGAGGAGGAGGAGGT |
| *CerS2* | GCGCAA AATGAGGTAGGC C | GCGCAAAATGAGGTAGGCC |
| *CerS3* | CATCCTGGATGCTCTTCATG | CGAGGTGATACATAGGCAAG |
| *CerS4* | CTTCGTGGCGGTCATCCTG | TGTAACAGCAGCACCAGAGAG |
| *CerS5* | AGAACGTGAGCTGGGCTGAT | GGCAATAAATCGCTCGAAGAG |
| *CerS6* | GCTTTGTTGCTGACGTGGACC | CCACTATGCTGGGTACGAGC |
| *Sgms1* | GTTCTCACCACAGTGATGATC | CAGTTGGGCTTCCCAGTCTC |
| *Sgms2* | GTGTGCTCCAAAGCTCAATGG | GTCAGTGTCAGCGTAACCGT |
| *Smpd2* | CTACCCACGGACCAGCAGA | CCCAGGCCAATCACATAGC |
| *Smpd3* | GCTTCAAGTGTCTCAACAGC | CCACCTGCACCTTGAGAAAC |
| *Smpd4* | GGCTGGGCTTTAGCTCCAT | CTTCGCTGAGCCGGAATATC |
| *Ugcg* | GAATGGCCGTCTTCGGGTTC | CACAAGAGAAGACACCTGGGAG |
| *GBA1* | GATACCAAGCTCAAGATACCC | GGTCTGGTGGTAGATGTCTC |
| *B4GALT6* | CCGGAACTATTACGGATGTGA | GTGTGCCAGTCTGTTCAATGA |
| *ELF1* | TGTTGTCCAACAGAACGACCT | GGAAAAATAGCTGGATCACCA |
| *ZFX* | GGATGATGCTGGCAAAATAGAAC | CAGTTCCACCTAAGTCATCTTC |
| *CTCF* | GACGAGTACCTGTGTGTGTG | CCAGTGTGAGCTTTGCAGTTA |
| *Actin* | ATTGGCAATGAGCGGTTCC | GGTAGAGTTTCGTGGATGCCACA |

**Supplementary Table S2.** List of Primers (human) used for validation of ChiP events.

| Gene Name | Forward primer (5' to 3') | Reverse primer (5' to 3') |
|---|---|---|
| *ZFX* | CTCTCCGGTCCCTTTTGATC | CGGAGCTTGTCTAGAAATCCA |
| *H3K4ME3* | CTCTCCGGTCCCTTTTGATC | CGGAGCTTGTCTAGAAATCCA |

**Supplementary Table S3** Table showing parameters for estimation of gangliosides from cell lines.

| Ganglioside Species | Precursor ion (Q1) | Product ion (Q3) | Collision Energy (V) |
|---|---|---|---|
| GM3-d3 | 1183.600 | 290.100 | -65.000 |
| d18:1/C16:0 GM1 | 1506.800 | 290.000 | -95.000 |
| d18:1/C16:0 GM3 | 1151.700 | 290.100 | -68.000 |
| d18:1/C18:0 GM3 | 1179.700 | 290.100 | -68.000 |
| d18:1/C20:0 GM3 | 1207.700 | 290.100 | -68.000 |
| d18:1/C22:0 GM3 | 1235.800 | 290.100 | -68.000 |
| d18:1/C23:0 GM3 | 1250.000 | 290.100 | -68.000 |
| d18:1/C24:0 GM3 | 1264.700 | 290.100 | -68.000 |
| d18:1/C16:0 GD3 | 721.200 | 290.200 | -55.000 |

#### 5. Legends for data sets (Data sets are in separate files)

**Data set 1.** Table showing the clinical and pathological information of Luminal A breast cancer female patients of Indian origin included in this study.

**Data set 2.** Table showing the immunohistochemistry data of tissue microarray from tumor samples of Indian, female breast cancer patients showing the score for UGCG and ZFX staining.

**Data set 3.** Original immunoblots and replicates used for quantification.
